# TTF1 control of LncRNA synthesis and cell growth delineates a tumour suppressor pathway acting directly on the ribosomal RNA Genes

**DOI:** 10.1101/2024.02.11.579707

**Authors:** Dany S. Sibai, Michel G. Tremblay, Frédéric Lessard, Christophe Tav, Marianne Sabourin-Félix, Mark D. Robinson, Tom Moss

**Affiliations:** St-Patrick Research Group in Basic Oncology, Cancer Division of the Quebec University Hospital Research Centre, Laval University, Quebec, Canada; Department of Molecular Biology, Medical Biochemistry and Pathology, Faculty of Medicine, Laval University, Quebec, Canada; Cancer Research Centre, Laval University, Quebec, Canada; Department of Molecular Life Sciences, University of Zurich, Zurich, Switzerland

**Author notes:** Correspondence should be addressed to; Tom Moss, PhD, Edifice St Patrick, 9 rue McMahon, Québec, QC, G1R 3S3, Canada. E-mail.

## Abstract

The tumour suppressor p14/19ARF regulates ribosomal RNA (rRNA) synthesis by controlling the nucleolar localization of Transcription Termination Factor 1 (TTF1). However, the role played by TTF1 in regulating the rRNA genes and in potentially controlling growth has remained unclear. We now show that TTF1 expression regulates cell growth by determining the cellular complement of ribosomes. Unexpectedly, it achieves this by acting as a “roadblock” to synthesis of the non-coding LncRNA and pRNA that we show are generated from the “Spacer Promoter” duplications present upstream of the 47S pre-rRNA promoter on the mouse and human ribosomal RNA genes. Unexpectedly, the endogenous generation of these non-coding RNAs does not induce CpG methylation or gene silencing. Rather, it acts *in cis* to suppress 47S preinitiation complex formation and hence *de novo* pre-rRNA synthesis by a mechanism reminiscent of promoter interference or occlusion. Taken together, our data delineate a pathway from p19ARF to cell growth suppression via the regulation of ribosome biogenesis by non-coding RNAs and validate a key cellular growth law in mammalian cells.

## INTRODUCTION

Tumour suppression and cellular senescence are related mechanisms that limit cell proliferation and the onset of cancer. The p14/p19ARF (ARF) tumour suppressor encoded by the CDKN2A locus has been implicated in both these mechanisms. The CDKN2A locus in fact encodes alternative reading frames for two distinct tumour suppressors, INK4A a CDK inhibitor controlling cell cycle progression and ARF a regulator of senescence and of p53 signalling. Both INK4A and ARF are very commonly deleted in human cancer, but the tight link of ARF inactivation with p53 inactivation suggested they have overlapping functions (Klimovich et al., 2019; Mina et al., 2017; Peters, 2008). The best-known of these is the ability of ARF to stabilize p53 by inhibiting the E3-ligase MDM2 and hence prevent its degradation. However, ARF also displays p53-independent activities both in tumour suppression and cellular senescence that are at least in part mediated by its ability to suppress ribosome biogenesis (Lessard et al., 2010; Sugimoto et al., 2003). The generation of ribosomes is an essential part of cell growth and indeed has been long known to bear a direct relationship with cell proliferation (Dai and Zhu, 2020; Scott et al., 2014). The higher rates of cell proliferation and growth associated with tumour growth should therefore require increased numbers of cellular ribosomes to cope with the enhanced need in protein synthesis. Thus, oncogenic transformation is unavoidably dependent on an ability to enhance ribosome biogenesis, while tumour suppression and cellular senescence are reliant on its suppression (Elhamamsy et al., 2022; Gaviraghi et al., 2019; Kusnadi et al., 2014). It is therefore not surprising that oncogenes and tumour suppressors invariably aim respectively to enhance or suppress ribosome biogenesis. The dependence of growth on ribosomes also makes ribosome biogenesis a logical target for chemotherapy, a subject that has stimulated recent interest in identifying inhibitory drugs (Drygin et al., 2010; Ferreira et al., 2020; Hein et al., 2013; Mars et al., 2020; Pelletier et al., 2018; Zisi et al., 2022).

The ribosome is the very large nucleoprotein complex responsible for cellular protein synthesis. Its biogenesis begins in the nucleolus with the synthesis of the ribosomal RNAs (rRNAs) and their co-transcriptional assembly into pre-ribosomal particles (Klinge and Woolford, 2019). The resulting mature ribosomes consist of 1/3^rd^ protein and 2/3rd rRNA and account for around 80% of total cellular RNA. Hence their synthesis represents a major metabolic task and one that limits cell proliferation by directly determining the cell’s ribosome complement and translational capacity. Transcription of the several hundred rRNA genes (the rDNA) is exclusively undertaken by RNA polymerase I (RPI/Pol1/PolR1) and is subject to regulation by a wide range of growth and nutrient signalling pathways as well as to dysregulation by oncogenes (Barna et al., 2008; Bywater et al., 2013; Moss et al., 2007). In recent years suppression of ribosome biogenesis has also been implicated in growth control as a major regulator of the p53 tumour suppressor (Pelletier et al., 2018). Thus, ribosome biogenesis is a central determinant of cellular growth.

As mentioned above, the ARF tumour suppressor has been implicated as a direct regulator of ribosomal RNA synthesis. ARF was initially found to regulate rRNA processing (Itahana et al., 2003; Sugimoto et al., 2003). Subsequently we found that ARF in fact inhibited rRNA synthesis and this was related to its ability to suppress the nucleolar localization of the RPI-specific Transcription Termination Factor 1 (TTF1) (Lessard et al., 2010). Nucleolar transport of TTF1 was shown to be mediated by the chaperone NPM1 but to depend on Nucleolar Localization Signals (NoLSs) within its short N-terminus repeat motifs (Boutin et al., 2019; Lessard et al., 2010; Lessard et al., 2012). Binding of ARF to these motifs was shown to suppress their NoLS activity and hence to delocalize TTF1 but how this resulted in the suppression of rRNA synthesis was at the time unclear. TTF1 is a Myb-related sequence-specific DNA binding factor and an orthologue of the yeast Reb1p “roadblock” termination factor (Colin et al., 2014). Like Reb1p, it terminates transcription of the primary rRNA precursor (47S pre-rRNA) by binding to multiple sites downstream of the rDNA coding region (Kuhn et al., 1990). But TTF1 also binds to sites upstream of the 47S pre-rRNA promoter where it has been shown to have contrasting and antagonistic activities of promoter activation by driving nucleosome displacement or rDNA silencing by catalyzing promoter methylation (Li et al., 2006; Santoro et al., 2002). Both these activities were found to be mediated by recruitment of the ACF/ISWI-related ATP-dependent remodelling complex NoRC (Li et al., 2006; Manelyte et al., 2014). NoRC recruitment to TTF1 and the induction of promoter methylation were shown to be enhanced by expression of a non-coding RNA (ncRNA) (Mayer et al., 2006; Schmitz et al., 2010). This promoter RNA (pRNA) is derived from a long non-coding RNA (LncRNA) complementary to the sense strand of the proximal rDNA Intergenic Spacer (IGS) and was thought to be transcribed from the Spacer Promoter duplications present in mouse, human and other higher eukaryote rDNAs (Mars et al., 2018; Moss et al., 2007; Savic et al., 2014). However, despite the activation-inhibition dichotomy of TTF1 function, little data exists on its regulation or that of the LncRNA and pRNAs, or their relationship with cell growth and senescence. Here we have investigated these questions and come to the surprising conclusions that in fact TTF1 acts as an *in cis* roadblock to Spacer Promoter driven LncRNA synthesis. We find that depletion of TTF1 enhances endogenous LncRNA and pRNA synthesis but does not cause rDNA methylation nor silencing. Rather, LncRNA synthesis destabilises the 47S preinitiation complex and suppresses 47S pre-rRNA synthesis by a mechanism reminiscent of promoter interference or occlusion. Most strikingly, TTF1 levels determine the rate of cell proliferation by directly regulating the cell’s ribosome complement. Natural accumulation of the p19ARF tumour suppressor during senescence also enhances LncRNA synthesis and reduces the cellular ribosome complement consistent with its role in regulating the nucleolar localization of TTF1. The data therefore delineate a previously unrecognized pathway of tumour suppression that acts via ribosome biogenesis to control cell growth.

## MATERIALS AND METHODS

### Generation of TTF1 conditional cell lines

Cell lines conditional for TTF1 were generated in two steps. A Doxycycline (Dox) regulated 3xFLAG-tagged TTF1 cDNA transgene was first introduced into NIH3T3 and cell clones selected for Dox-dependent FLAG-TTF1 expression. The endogenous *Ttf1* alleles were then inactivated using CRISPR/Cas9 and a guide RNA targeting exon 2, a silent mutation in the 3xFLAG-TTF1 cDNA preventing its CRISPR/Cas9 targeting (Figure S1A). Viable clones were initially selected in media containing 10ng.ml^-1^ Dox and screened for inactivation of both endogenous *ttf1* alleles Figure S2. Two such clones (clones C8 and C14) displayed strong Dox-dependent expression of Flag-TTF1 (Figure S1B). TTF1 expression was reproducibly induced to near endogenous levels (50 to 60% of NIH3T3) at 100ng.ml^-1^ Dox in clone C8 and at 10ng.ml^-1^ in clone C14 (Figure S1C). Concomitantly, background expression of TTF1 in the absence of Dox was consistently very low (∼2% endogenous) in clone C8 but tended to be a little higher and more variable in clone C14 (Figure S1C). All cell lines were cultured at 37 deg.C and 5% CO_2_ in high glucose Dulbecco’s Modified Eagle’s Medium (DMEM), 2.2g.l^-1^ NaHCO_3_ supplemented with 10% fetal bovine serum (FBS), penicillin/streptomycin (Wisent) and 2 mM L-glutamine (Wisent).

### Primary antibodies for ChIP and Western blotting

Rabbit polyclonal antibodies against UBF, N-terminal and C-terminal domain of TTF1, RPI large subunit (RPA194/POLR1A), TAF1B were generated in the laboratory and have been previously described (Herdman et al., 2017). All other antibodies were obtained commercially; anti-FLAG M2 (F3165, Sigma), anti-Tubulin (Cell Signaling #38735), anti-GAPDH (MediMabs #MM-0163-P), p19ARF (Santa Cruz Biotechnology #sc-32748).

### Western Blotting

TTF1 protein levels were monitored by Western blotting. At harvesting, cells were quickly rinsed in cold phosphate buffered saline (PBS), recovered by centrifugation (2 min, 2000 r.p.m.) and resuspended directly in SDS–polyacrylamide gel electrophoresis (SDS-PAGE) loading buffer (Laemmli, 1970). After fractionation by SDS-PAGE, proteins were analysed by standard Western blotting procedures using an HRP conjugated secondary antibody and Immobilon chemiluminescence substrate (Millipore-Sigma). Membranes were imaged on an Chemidoc^TM^ MP (Biorad) gel imaging system and TTF1 levels were determined from lane scans using ImageJ and/or ImageQuant (Cytiva).

### Cell Proliferation Assay

Cells were continuously cultured for more than a week prior to assay. Cells were plated at ∼500 per well in 96-well plates and cultured for five days. At each timepoint, duplicate wells were treated with Hoechst 3342 (ThermoFisher Scientific) for 45min. Images were acquired using a Cytation5 (BioTek) and cell counts for each cell line were determined using the Gen5 software.

### Fluorescence-activated cell sorting (FACS)

Clone 8 and Clone 14 were cultured in the presence and absence of Dox as indicated above. EdU (5-ethynyl 2′-deoxyuridine) was added to the culture medium to a final concentration of 10 μM and labelling allowed to proceed for 2 hours. Cells were then trypsinized, counted, resuspended in PBS, 1% BSA and fixed and labelled using the Click-iT Plus EdU Alexa Fluor 488 Flow Cytometry Kit and the FxCycle™ Far Red cell cycle stain (Invitrogen, ThermFisher Scientific). The cell suspension was then passed through a pre-separation filter to remove cell aggregates and analyzed on a BD-FACS Aria II flow cytometer (Beckton Dickinson, Franklin Lakes, NJ). FACS data was then analyzed using the FlowJo (Beckton Dickinson) and FCS-alyzer (Sven Mostböck) software. The unlabelled control cells were also analyzed to assess the minimal fluorescein isothiocyanate (FITC) baseline.

### Total RNA Extraction and Quantification, cDNA synthesis and RT-PCR

Cells were trypsinized, counted and total RNA was recovered from 3×10^6^ cells using 1 ml of Trizol (ThermoFisher Scientific) according to the manufacturer’s protocol. RNA yields were determined using the Qubit RNA BR assay kit (ThermoFisher Scientific). cDNA was synthetized from 2ug of total RNA using MMLV-Reverse transcriptase and d(N)_6_ random primers in a 20ul reaction following the manufacturer’s protocol (ThermoFisher Scientific). RT-qPCR analyses were performed in triplicate using 2.5 μl of cDNA, 20 pM of each primer, and 10 μl of Quantifast SYBR Green PCR Master Mix (QIAGEN) or PowerUp™ SYBR™ Green Master Mix (ThermoFisher) in a final volume of 20 ul. Forty reaction cycles of 15 s at 95 ^0^C, 30 s at 58 ^0^C and 30 s at 72 ^0^C were carried out on a Multiplex 3005 Plus (Stratagene/Agilent). The amplicon coordinates relative to the 47S rRNA initiation site (BK000964v3) were as follows: Tsp, 43253 to 43447; Tsp4, 43316 to 43404; Prom2, 45144 to 45308; T2440, 15868 to 15976.

### Determination of rRNA synthesis rate

The rate of rRNA synthesis was determined by metabolic labelling immediately before cell harvesting. 10 µCi [^3^H]-uridine (PerkinElmer) was added per 1ml of medium and cell cultures incubated for a further 30min to 7h as indicated. RNA was recovered with 1 ml Trizol (ThermoFisher Scientific) according to the manufacturer’s protocol and resuspended in Formamide (ThermoFisher Scientific). One microgram of RNA was loaded onto a 1% formaldehyde/MOPS Buffer gel (Stefanovsky and Moss, 2016; Stefanovsky et al., 2001) or a 1% formaldehyde/TT Buffer gel (Mansour and Pestov, 2013). The EtBr-stained gels were photographed using the G:BOX acquisition system (Syngene), irradiated in a UV cross-linker (Hoefer) for 5 min at maximum energy, and transferred to a Biodyne B membrane (Pall). The membrane was UV cross-linked at 70 J/cm^2^, washed in water, air dried and exposed to a Phosphor BAS-IP TR 2025 E Tritium Screen (Cytiva). The screen was then analyzed using a Typhoon imager (Cytiva) and quantified using the ImageQuant TL image analysis software.

### Psoralen crosslinking accessibility and Southern blotting

The psoralen crosslinking accessibility assay was performed on isolated cell nuclei as previously described (Sanij et al., 2008; Stefanovsky and Moss, 2006) and DNA probed using the ^32^P-labelled 6.7kb rDNA EcoRI fragment (pMr100) (Conconi et al., 1989; Grozdanov et al., 2003). The ratio of “active” to “inactive” genes was estimated by analyzing the intensity profile of low and high mobility bands revealed by phospho-imaging on an Amersham Typhoon (Cytiva) using ImageQuant TL.

### Estimates of rDNA CpG methylation

rDNA Copy Number and CpG methylation were determined using both a Southern blot technique and by WGBS/EM-Seq. For Southern blot analysis, 2µg of total DNA was digested with BamHI, BamHI+XmaI or BamHI+SmaI and fractionated by gel electrophoresis and processed by standard blotting procedures. The EtBr-stained gels were photographed (G:BOX, Syngene), DNA sheared by UV irradiation (5 min in Hoefer UVC500 at maximum energy) and transferred to a Biodyne B membrane (Pall). The membrane was UV cross-linked at 70 J/cm^2^ (Hoefer UVC500) washed in water and co-hybridized with a ^32^P-labelled 1.02 kb PflMI-BamHI fragment from the 28S gene and a 0.78 kb BamHI-EcoRI covering exons 15 and 16 of the *Ubtf* gene as single copy internal standard. Bands were revealed by phospho-imaging using an Amersham Typhoon (Cytiva) and quantified using the ImageQuant TL software. The use of the WGBS/EM-Seq was also performed on total DNA using the NEBNext MethylSeq library protocol and sequencing on Illumina NovaSeq 6000 (McGill University and Genome Quebec Innovation Centre). Sequences were aligned and analyzed using Bowtie 2 (Langmead and Salzberg, 2012) and Bismark 0.15 (Krueger and Andrews, 2011).

### RDNA copy number estimation

Relative rDNA copy numbers were estimated from Southern blots data as the intensity ratio of the BamHI 4.8 kb rDNA fragment over the 7.9 kb single copy *Ubtf* BamHI fragment and normalized to this ratio for the wild type NIH3T3 (Figure 4E and S6B to D). rDNA copy numbers were also determined from whole genome sequencing as the ratio of the number of reads aligned to the rDNA over the total of number of aligned reads, and again normalized to the equivalent data for wild type NIH3T3 (Figure S6C).

### LncRNA transcript mapping by S1 Nuclease protection

Total RNA was harvested using 1 ml Trizol (ThermoFisher Scientific) according to the manufacturer’s protocol and resuspended in Formamide (ThermoFisher Scientific). 24 ul (5ug) of RNA, 3ul of the 5’ ^32^P labelled rDNA probe (Figure S8B) and 3 ul of 10x S1 hybridization buffer (4M NaCl, 0.4M Na-PIPES pH6.4) were combined, denatured at 100 deg.C for 1 min. and then incubated overnight at 56°C. 100U of S1 Nuclease (ThermoFisher Scientific) in 270 ul of ice-cold S1 reaction buffer (30mM Na actetate, 2mM ZnSO_4_, 0.2M NaCl, pH4.5) was added followed by incubation for 1h at 37°C, and nucleic acids recovered by phenol/chloroform extraction and EtOH precipitation. The resulting protected fragments were fractionated by electrophoresis on an 8% Tris-Borate-7M Urea gel, visualized by exposure to a Phosphor BAS-IP screen (Cytiva) and subsequent imaging on an Amersham Typhoon (Cytiva), and quantified using the ImageQuant TL software.

### Chromatin immunoprecipitation (ChIP)

Cells were fixed with 1% formaldehyde for 8 min at room temperature. Formaldehyde was quenched by addition of 125 mM Glycine and cells harvested and washed in PBS. Nuclei were isolated using an ultrasound-based nuclei extraction method (NEXSON: Nuclei Extraction by SONication) (Arrigoni et al., 2016; Herdman et al., 2017; Tremblay et al., 2022) with some modifications. Briefly, for all cell types, 33 million cells were resuspended in 1.5 ml of Farnham buffer (5 mM PIPES pH 8.0, 85 mM KCl, 0.5% IGEPAL, 1mM PMSF and 1ug.ml^-1^ each of Aprotinin, Leupeptin, Pepstatin). Cell suspensions were sonicated in 15 ml polystyrene tubes (BD #352095) using 3 to 4 cycles of 15 sec on : 30 sec off at low intensity in a Bioruptor (Diagenode). After recovery of the NEXSON-isolated nuclei by centrifugation (1000g, 5 min), nuclei were resuspended in 1.5 ml of shearing buffer (10 mM Tris-HCl pH 8.0, 1 mM NaEDTA, 0.1% SDS and protease inhibitors) and sonicated for 25 min, 30 sec on : 30 sec off, at high intensity. Each immunoprecipitation (IP) was carried out using the equivalent of 16 x 10^6^ cells as previously described (Herdman et al., 2017).

### ChIP-qPCR analysis

All ChIP experiments included a minimum of 3 biological replicates and were analyzed as previously described (Herdman et al., 2017). For qPCR analysis, reactions (20 μl) were performed in triplicate using 2.5 μl of sample DNA, 20 pmol of each primer, and 10 μl of Quantifast SYBR Green PCR Master Mix (QIAGEN) or PowerUp™ SYBR™ Green Master Mix (ThermoFisher). Forty reaction cycles of 15 s at 95 °C, 30 s at 58 °C and 30 s at 72 °C were carried out on a Multiplex 3005 Plus (Stratagene/Agilent). The amplicon coordinates relative to the 47S rRNA initiation site (BK000964v3) were as follows: IGS3, 42653-42909; SpPr, 43076-43279; Tsp, 43253-43447; Tsp4, 43316 to 43404; T0/Pr (47SPr), 45133-40; 47S-5’, 159-320; ETS, 3078-3221; 28S, 10215-10411; T1-3, 13417-13607. Data was analyzed using the MxPro software (Agilent). The relative occupancy of each factor was determined by comparison with a standard curve of amplification efficiency for each amplicon using a range of input DNA amounts generated in parallel with each qPCR run.

### ChIP-Seq and data analysis

ChIP DNA samples were quality controlled by qPCR before being sent for library preparation and 50 base single-end or 100 base paired-end sequencing on an Illumina HiSeq 2500 or 4000, or NovaSeq 6000 (McGill University and Genome Quebec Innovation Centre). Sequence alignment and deconvolution of factor binding profiles to remove sequencing biases (Deconvolution ChIP-Seq, DChIP-Seq) were carried out as previously described (Herdman et al., 2017; Mars et al., 2018).

### RNA-Seq and data analysis

Clone C8 was cultured for >7days with 100ng.ml^-1^ doxycycline (Dox100), cells were then either harvested or cultures continued for a further 3, 5 or 7 days without doxycycline (Dox0). Cultures were then either harvested or 100ng.ml^-1^ doxycycline readded and cultures maintained for a further 3, 5, and 7 days (Dox100-Readdition). Total RNA was extracted from these cultures and from parallel cultures of NIH3T3 using Trizol and GeneJet RNA micro kit (ThermoFisher Scientific). RNA sequencing libraries were created using the NEBNext mRNA stranded protocol (New England Biolabs) and 100 base paired-end sequencing was performed on an Illumina NovaSeq 6000 (McGill University and Genome Quebec Innovation Centre). Raw sequencing reads were subjected to quality control using FastQC (version 0.11.9) and were processed using the MUGQIC RNA-Seq pipeline (version 4.4.2) (Bourgey et al., 2019). In brief, reads were trimmed for adaptor sequences using Trimmomatic (Bolger et al., 2014). High-quality reads were aligned to the mouse reference genome (GRCm38) using the STAR aligner (version 2.7.8a) (Dobin et al., 2013). Gene counts were determined using featureCounts (version 2.0.3) (Liao et al., 2014) with the genomic annotation Ensembl release 102. Differential expression analysis was performed using the edgeR package (version 3.40.2) (Robinson et al., 2010) in R (version 4.2.2). Normalization factors were calculated using the calcNormFactors function with the default ‘TMM’ method. Dispersion was estimated with the estimateDisp function. Differential expression was tested using the Genewise Negative Binomial Generalized Linear Models with Quasi-likelihood Tests (glmQLFit) provided by edgeR. A gene was considered significantly differentially expressed if p-value, adjusted for False Discovery Rate (FDR), was less than or equal to 0.05. Visualization of differential gene expression analysis was performed using the EnhancedVolcano (version 1.16.0) R package.

## RESULTS

Our previous DChIP-Seq analyses created a detailed map of the mouse and human rDNA chromatin and the recruitment of RPI and its basal factors to the 47S Promoter, the enhancer repeats and the upstream Spacer Promoter (Herdman et al., 2017; Mars et al., 2018). These maps provided support for the notion that the rDNA LncRNA was generated by transcription from the Spacer Promoter. However, they also suggested that if this were the case, LncRNA synthesis would be prematurely terminated by TTF1 bound at a proximal site.

### Arrest of transcription from the Spacer Promoter

DChIP-Seq mapping previously showed that in mouse the Spacer and 47S promoters recruited SL1 and UBTF with equal avidity, and both recruited RPI polymerase (Herdman et al., 2017; Mars et al., 2018; Tremblay et al., 2022). However, unlike the recruitment of RPI into functional elongation complexes at the 47S promoter, RPI accumulated in a narrow binding peak immediately adjacent to the Spacer Promoter (Figure 1A). DChIP mapping revealed that this peak was centered just 30 b.p. downstream of the Spacer Promoter initiation site and hence represented very early RPI elongation complexes before completing promoter release. These immediately abutted the RPI specific Transcription Termination Factor 1 (TTF1) bound at a canonical site centred just 38 b.p. further downstream (Tsp in Figure 1A and 1B). This arrangement was also conserved in both the human and rat rDNAs, where we could identify TTF1 binding sites lying 64 b.p. and 66 b.p. downstream of the respective Spacer Promoters initiation sites (Mars et al., 2018; Smith et al., 1990). Inhibition of RPI transcription initiation by CX-5461(Mars et al., 2020) was observed to slightly skew the profile of “arrested” RPI upstream towards the Spacer Promoter initiation site and away from TTF1 as might be expected for arrested elongation complexes (Figure 1B). These data therefore inferred that TTF1 controlled transcription from the Spacer Promoter and hence most probably synthesis of both the LncRNA and pRNA. To directly test this possibility, we generated cell lines in which TTF1 levels could be regulated.

**Figure 1.**
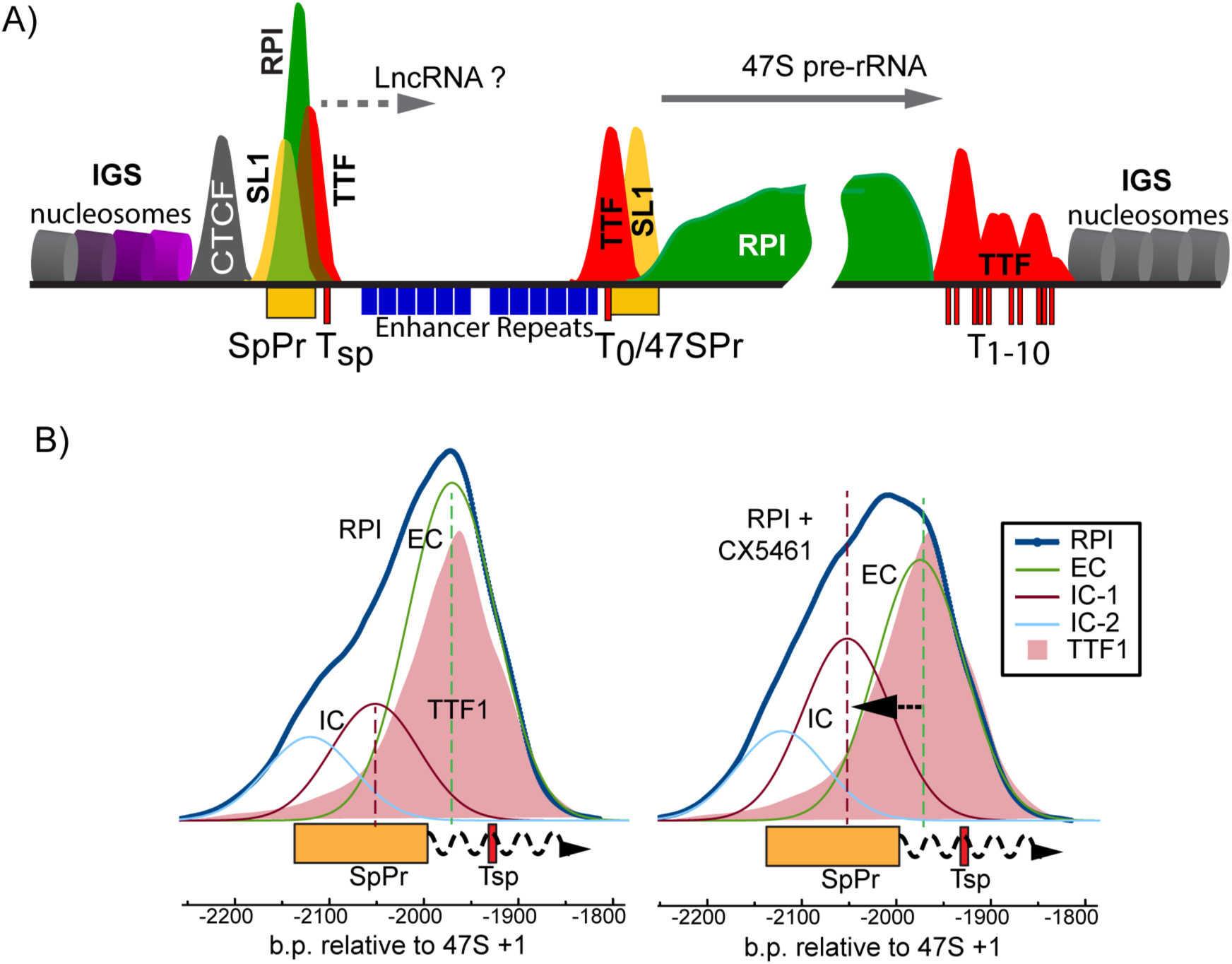
Organisation of the rDNA chromatin in mouse. A) Diagrammatic representation of mapping data for the major RPI transcription factors showing the Spacer (SpPr) and 47S (47SPr) Promoters, the canonical TTF1 binding sites T_sp_, T_0_, T_1-10_ and typical occupancy profiles for RPI, TTF1, the TBP-TAF complex SL1 and CTCF. B) DChIP mapping data for RPI and TTF1 at the Spacer Promoter before and after inhibition of RPI transcription initiation with CX-5461 (https://www.ebi.ac.uk/arrayexpress/experiments/E-MTAB-9242). The mapping profile plots for RPI are shown outlined in blue and for TTF1 in solid shading. Curve fits to the RPI profile are also shown and labelled EC for potential elongation, and IC for initiation complexes. The map positions of the Spacer Promoter and canonical TTF1 binding site are indicated below each plot along with the base positions relative to the 47S pre-rRNA initiation site as +1.

### Generation of TTF1 conditional cell lines

Since TTF1 loss was likely to be lethal (Lessard et al., 2010; Lessard et al., 2012), a doxycycline (Dox) regulated 3xFLAG-tagged TTF1 cDNA transgene modified to be resistant to subsequent CRISPR/Cas9 targeting was introduced into NIH3T3 Mouse Embryonic Fibroblasts (MEFs) and both endogenous *Ttf1* alleles then inactivated (Figure S1A and S2). The NIH3T3 cell line was chosen as starting point since it has been extensively studied and lacks a functional p19ARF gene whose expression might otherwise interfere with rDNA activity and TTF1 subcellular localization (Lessard et al., 2010; Sugimoto et al., 2003). The resulting cloned lines C8 and C14 displayed Dox-inducible expression of the exogenous 3xFLAG-TTF1 and no detectable expression of endogenous TTF1 (Figures 2A and S1B). At optimal Dox concentration, 3xFLAG-TTF1 levels in these cell lines approached those observed for endogenous TTF1 in the parent NIH3T3 cell line, respectively 0.8 ± 0.1 for C8 at 100ng.ml^-1^ and 0.6 ± 0.07 for C14 at 10ng.ml^-1^ Dox (respectively Dox100 and Dox10 in Figure 2A and S1C). The C8 cell line also displayed extremely low basal levels of 3xFLAG-TTF1 in the absence of Dox, (0.03 ± 0.005 relative to TTF1 in NIH3T3), while basal levels in the C14 line were some 3 times higher (0.1 ± 0.02) (Figures 2A and S1B&C). Chromatin Immunoprecipitation (ChIP) further confirmed the Dox dependent recruitment of the exogenous 3xFLAG-TTF1 to the canonical Tsp, T0 and T1-3 binding sites in both conditional cell lines (Figure S1D).

**Figure 2.**
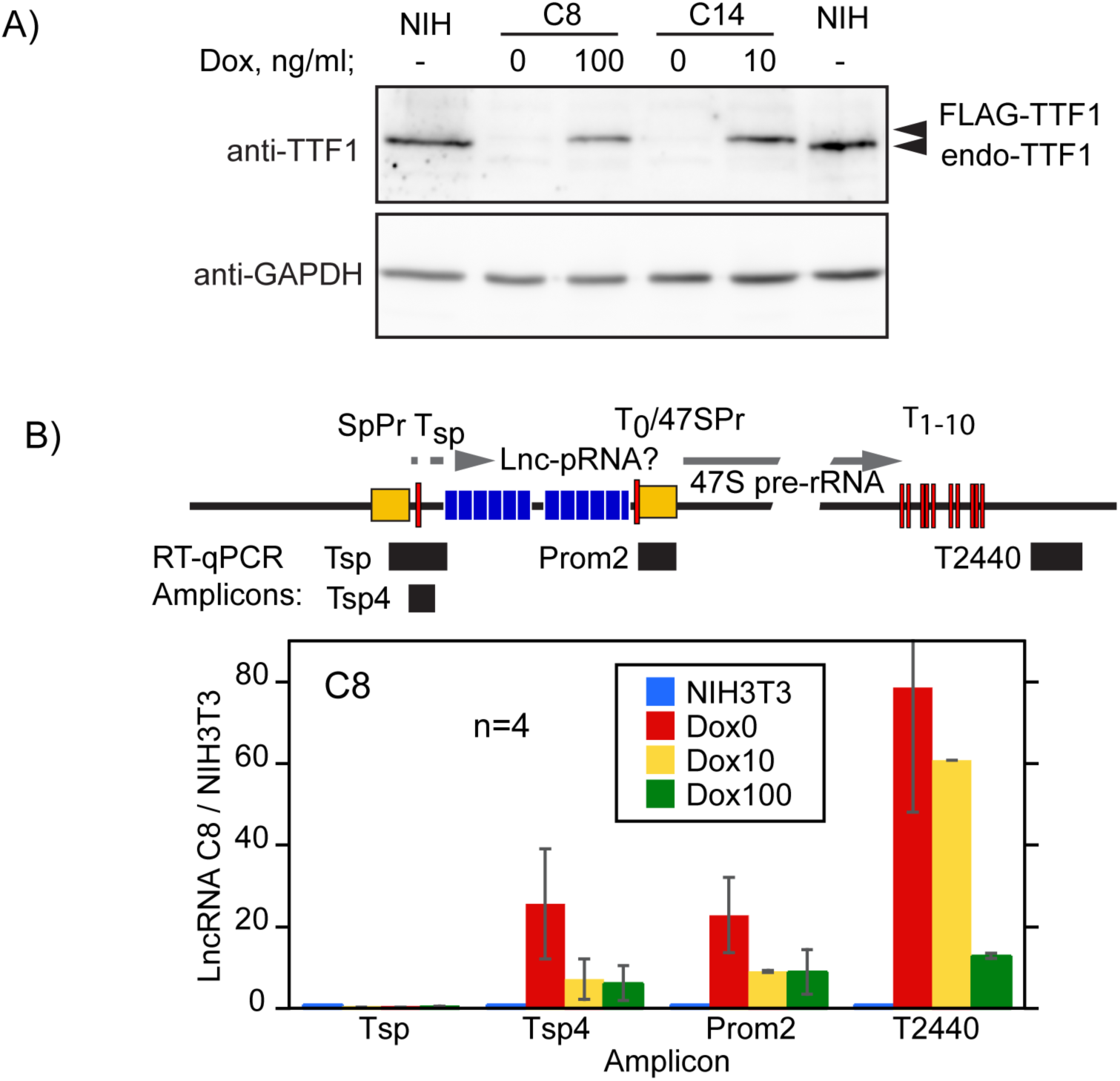
TTF1 regulates LncRNA synthesis. A) Expression of exogenous and endogenous TTF1 in clones 8 and 14 and in wild type NIH3T3 determined by western blotting in comparison to GAPDH levels. B) Upper panel; organisation of the mouse rDNA locus indicating the positions of the Spacer (SpPr) and 47S (47S Pr) promoter sequences and the canonical TTF1 binding sites Tsp, T_0_ and T_1-10_. The positions of the RT-qPCR amplicons are indicated. Lower panel; RT-qPCR analysis of RNA from clone C8 grown under the indicated conditions of TTF1 induction (Dox0, 10 and 100 refer respectively to growth with 0, 10 or 100ng.ml^-1^ Dox). Data were normalized to RPS12 mRNA levels (Savic et al., 2014) and are presented relative to wild type NIH3T3. The data derive from 4 independent biological replicas (n = 4) each analyzed in triplicate, and error bars indicate the SEM.

### TTF1 regulates LncRNA synthesis

Since we hypothesized that TTF depletion should prevent the premature arrest of Spacer Promoter transcripts and enhance LncRNA synthesis, we used RT-qPCR to detect transcripts downstream of the Spacer Promoter proximal TTF1 binding site Tsp (Figure 2B and S3). In the absence of Dox (Dox0), RT-qPCR detected transcripts crossing the Tsp site and the 47S Promoter (amplicons Tsp4 and Prom2) in both C8 and C14 cell lines (Figures 2B and S3). These transcripts were absent from the NIH3T3 parent cell line and were clearly derived from the Spacer Promoter since no RNA was detected crossing its initiation site (Tsp amplicon). As expected, depletion of TTF1 also allowed readthrough of the 47S pre-rRNA termination sites (T_1-10_) (amplicon T2440). In contrast, expression of exogenous FLAG-TTF1 in C8 cells maintained at the optimal 100ng.ml^-1^ Dox concentration (Dox100) significantly though not fully suppressed the LncRNA and pRNA transcripts and 47S readthrough, while the intermediate level of TTF1 at 10ng.ml^-1^ Dox (Dox10) provided less effective suppression of these RNAs. Similar results were obtained in C14 cells, except that in the absence of Dox, LncRNA, pRNA and 47S readthrough transcripts were 4 to 5 times lower than in C8 cells (Figures 2B and S3), and hence consistent with the higher basal TTF1 expression in these cells (Figure S1C).

### Depletion of TTF1 and enhanced LncRNA synthesis anti-correlate with cell growth rate

The proliferation rate of the TTF1 conditional cell lines displayed a strong dependence on the level of Dox induction. In the absence of Dox (Dox0), C8 cell cultures grew slowly with a doubling time of 84 h, as compared to 52 h at suboptimal (Dox10) and 38 h at optimal (Dox100) TTF1 transgene induction (Figure 3A and S4A). The slow-growth phenotype at Dox0 corresponded with a small (∼11%) increase in G1-phase cells at the expense of the S- and G2-phase cell populations, suggesting a slowing of progression through the G1/S boundary (Figure 3B). Strikingly, both the cell-cycle and the proliferation rate changes were reversed by re-induction of TTF1 transgene expression (Dox100Re in Figure3B and S4D). At Dox0 the C14 cells also showed a reduced doubling time of 57 h as compared to 32 h at optimal (Dox10) transgene induction (Figure S4B and C) and a minor 3 to 4% increase in the G1-phase cell population. Thus, depletion of TTF1 and expression of the LncRNA corresponded with a reduction in cell proliferation and a somewhat slowed entry into S-phase but both changes were rescued by re-induction of TTF1.

**Figure 3.**
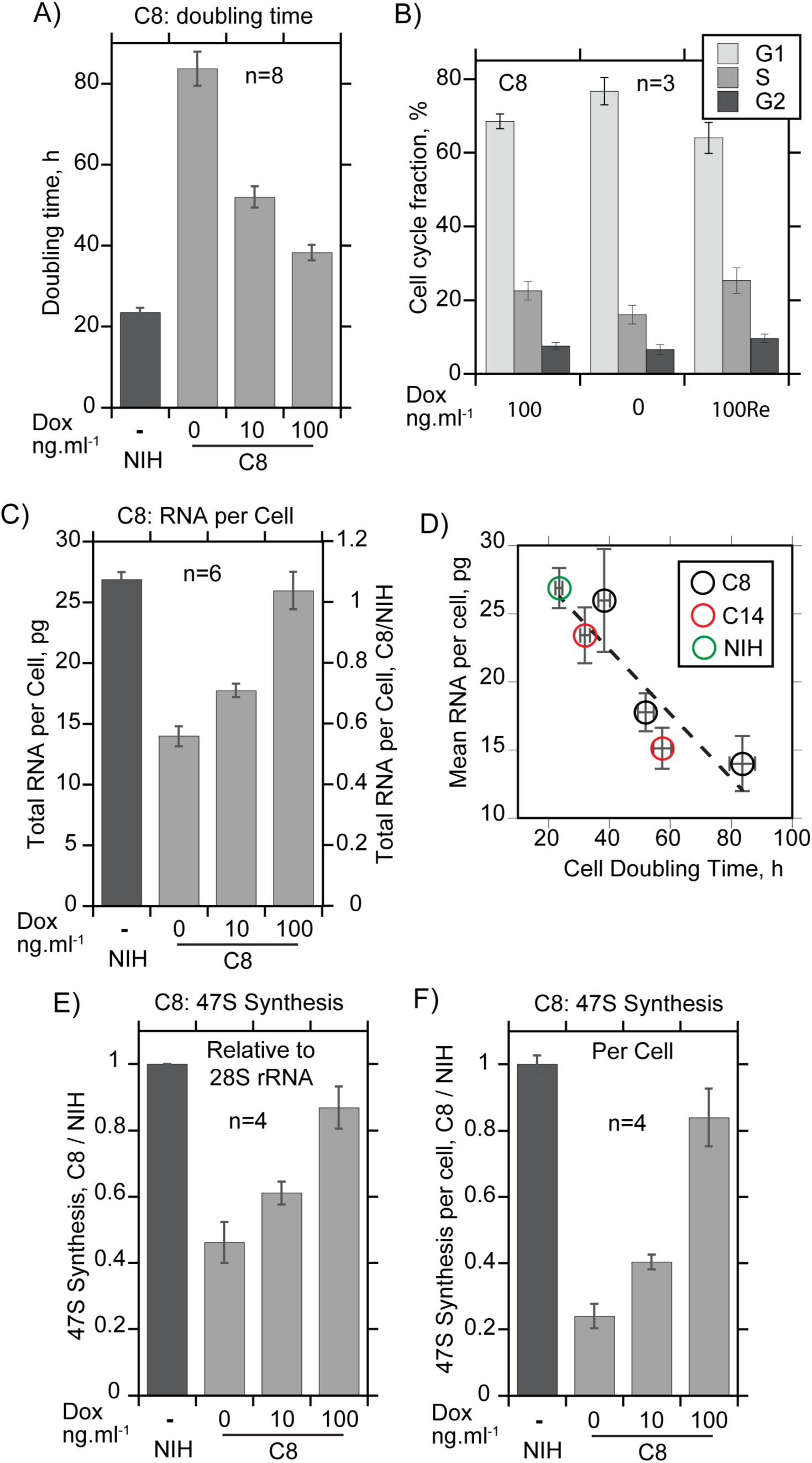
TTF1 determines cell proliferation and the cellular ribosome complement. A) The cell doubling time for clone C8 under different conditions of TTF1 induction was measured respectively as 84h, 52h and 38h (Dox0, Dox10, Dox100) and as 23h for NIH3T3. B) Cell cycle distribution of the C8 cells maintained optimally for ≥7 days in Dox100 or after 7 days TTF1 depletion in Dox0 followed by ≥7 days TTF1 reinduction (Doc100Re). C) Total RNA content of the clone C8 and of NIH3T3 cells grown under the same conditions as in A. D) Relationship between mean total RNA per cell (proportional to the cellular ribosome complement) and the cell doubling time taken from the data in A and B, and Figure S4C and E. E) and F) 47S pre-rRNA synthesis in clone C8 and NIH3T3 grown as in A as determined by [^3^H]-uridine metabolic pulse labeling (7 h). Panel E shows the data normalized to total 28S rRNA and Panel F to total RNA per cell. See Figure S4A for growth curve data and Figure S4B, C, E and F for equivalent datasets for clone C14. “n” indicates the number of biological replicas and error bars the SEM.

### TTF1 determines the cellular ribosome complement

Strikingly, depletion of TTF1 led to a strong reduction in total cellular RNA. Fully TTF1 depleted C8 cells contained half as much total RNA as the same cells grown under conditions of optimal TTF1 induction (Dox100), while at Dox10 they contained 30% less RNA (Figure 3C). C14 cells also contained 30 to 35% less RNA when grown at Dox0 as compared to growth at near optimal TTF1 expression (Dox10) (Figure S4E). In neither case could these very significant RNA deficits be explained by the observed small change in cell cycle distribution, which would in the most extreme case (C8 cells at Dox0) account for only a 6% reduction in total RNA.

Since total cellular RNA is constituted of ∼80% ribosomal RNA, its reduction on TTF1 depletion represented a very significant deficit in the ribosome complement of the cells, and this provided a rational explanation for the slow-growth phenotype. A central axiom of the so-called cell growth laws is the inverse relationship between the cellular ribosome complement and the cell doubling time (Dai and Zhu, 2020; Scott and Hwa, 2011; Scott et al., 2014). Analysis of the data for the C8 and C14 cell lines showed that this relationship was valid over the different conditions of TTF1 expression (Figure 3D). Thus, TTF1 levels appeared to determine the cellular proliferation rate by regulating the cellular ribosome complement.

### TTF1 controls the cellular ribosome complement by determining the rate of rRNA synthesis

The ribosome complement of a cell is predominantly determined by the balance between the rate of ribosome synthesis and its dilution by cell division. Since rRNA synthesis is the primary determinant of ribosome production, we used metabolic RNA labelling to determine its rate in the TTF1 conditional cells (Figure 3E and S4F). When normalized to mature steady-state rRNA, *de novo* synthesis of 47S pre-rRNA in the C8 cells ranged from 46% at Dox0 to 87% at Dox100 of that in NIH3T3. Since the cellular concentration of the mature rRNAs also varied with the growth conditions, this meant that the actual per cell synthesis rates were respectively 22% at Dox0 and 82% at Dox100 of NIH3T3 (Figure 3E and F). 47S pre-rRNA synthesis in the C14 cells displayed a similar but narrower range of per cell synthesis rates (Figure S4F), consistent with the smaller range of TTF1 regulation in these cells. Thus, the reduced rRNA synthesis rates and extended cell doubling times together explained the changes in the cellular ribosome complement observed on TTF1 depletion. For example, C8 cell cultures maintained at Dox0 took 3.5 times longer to divide and synthesized rRNA at 0.22 times the rate of NIH3T3 cultures (Figure 3A & F). They would then be expected to generate 0.22 x 3.5 = 0.77 ± 0.1 of the ribosome complement of NIH3T3 during each cell cycle. Though this is somewhat higher than the measured value of 0.52 ± 0.05 (Figure 3C), it assumes that all newly synthesized rRNA was stably integrated into ribosomes despite the obvious stresses of TTF depletion and slow growth. Hence, the data strongly suggested that TTF1 levels determined the cellular ribosome complement and growth rate by regulating the rate of rRNA synthesis. However, this left open the question of whether TTF1 levels mediated this effect directly at the level of the rDNA or indirectly via the regulation of other genes.

### TTF1 depletion has no effect on the expression of genes other than the rDNA

TTF1 is a sequence-specific DNA binding factor and distant relative of the Myb transcription factor family but to date it has only been implicated in rRNA gene expression. However, to test this we performed an RNA-Seq analysis before and after TTF1 depletion. The data revealed that the strong TTF1 depletion observed in C8 cells at Dox0 had no detectable effect on the expression of genes other than the rDNA and excluded major effects of TTF1 beyond the rDNA (Figure 4A and S5). For example, using fold change (FC) cut-offs of log_2_ > 1 and log_2_ < −1 and an adjusted p-value (FDR) of < 0.05, no significant change in mRNA gene expression was detected either after strong TTF1 depletion (e.g., Dox100 Day0 control versus Dox0 Day5 depletion), or even after strong depletion followed by TTF1 reinduction (e.g., Dox100, Day0 control versus Dox100 re-addition, Day5). Thus, the expression level of TTF1 determined the cellular ribosome complement and cellular growth exclusively by regulating the rRNA genes.

**Figure 4.**
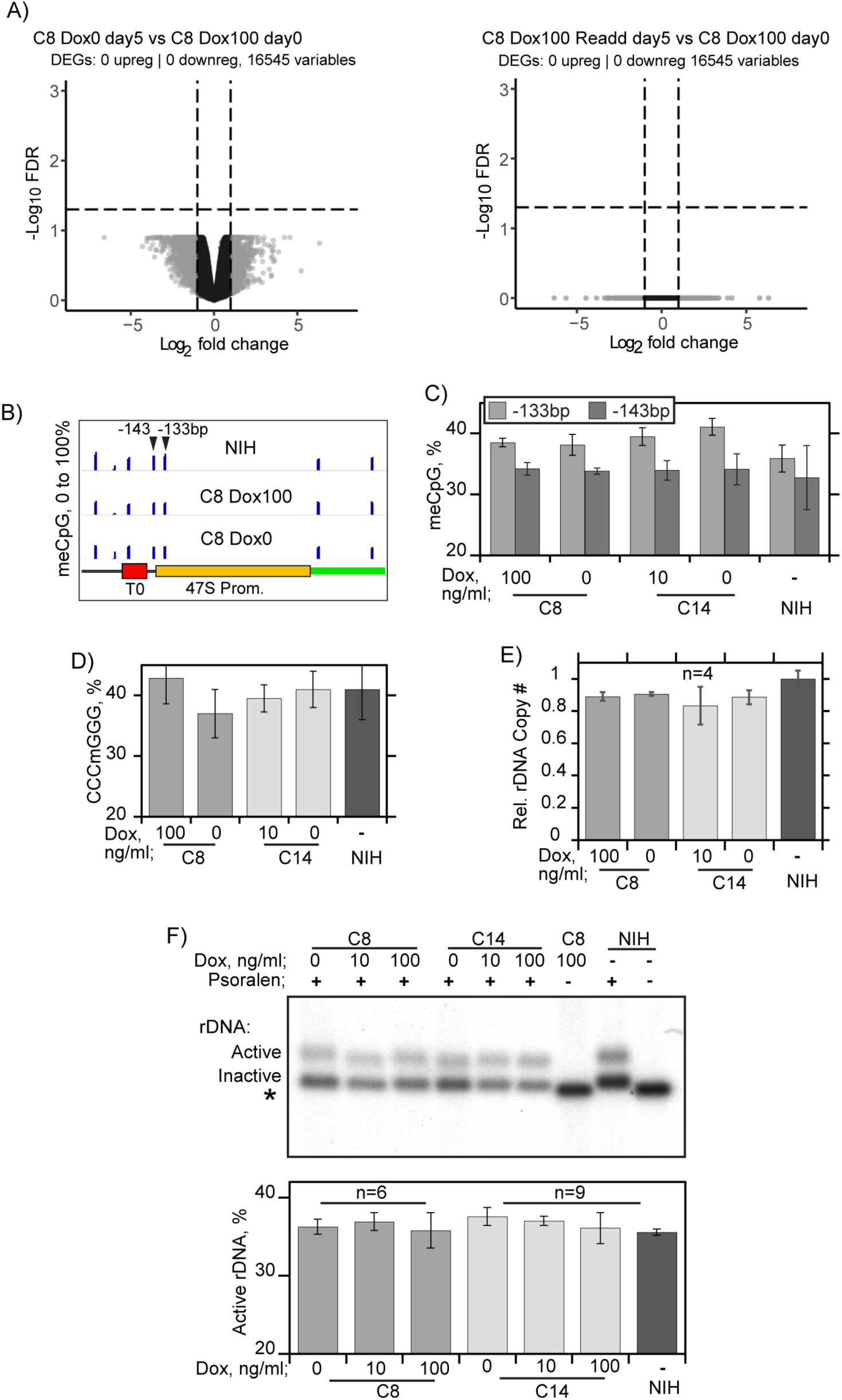
TTF1 depletion does not affect gene expression genome-wide, nor does it enhance rDNA methylation or silencing. A) RNA-Seq analyses of TTF1 depletion and reinduction conditional clone C8. Panels show plots of False Discovery Rate (FDR) against Fold Change for mRNA expression in biological duplicate RNA samples. Left panel, C8 cultures grown for 5 days without TTF1 induction (Dox0), or Right panel, 5 days after readdition, each compared to the starting (Dox100) culture. Using an adjusted p-value of less than 0.05, no significant change in mRNA expression was detected due TTF1 depletion (Dox0 vs Dox100) or between starting cell cultures and those subjected to TTF1 depletion and readdition (Dox100-Readd vs Dox100). Corresponding TTF1 expression was monitored by Western blot, (Figure S5A). B) Detailed view and C) quantification of critical 47S promoter CpG methylation sites in NIH3T3 and in clones C8 and C14 maintained at minimal (Dox0) or optimal (Dox 100/10) TTF1 expression levels as determined by WGB-Seq/EM-Seq, see also Figure S6A. D) CpG methylation of 47S rRNA coding sequences estimated by inhibition of cleavage of CCCmGGG (SmaI) sites is also not affected by TTF1 depletion, see Figure S6B for detail. E) TTF1 depletion also does not significantly affect rDNA copy number. The mean clone C8 and 14 rDNA copy numbers from whole genome sequencing and Southern blot analyses are shown relative to those for NIH3T3, see Figure S6B to D for more detail. F) Psoralen Accessibility Crosslinking (PAC) reveals no significant effect of TTF1 depletion on the levels of rDNA silencing, which remained constant throughout. The upper panel shows an example of PAC analysis for NIH3T3 and clones C8 and C14 maintained at various levels of Dox. The lower panel shows the mean “Active” rDNA fraction determined from PAC analyses of the 1.3, 2.4 and 4.7 kbp rDNA BamHI fragments. “n” indicates the number of biological replicas and error bars the SEM.

### TTF1 regulation of rRNA synthesis does not involve CpG methylation, rDNA silencing or rDNA copy number change

Given that TTF1 expression determined the cell growth rate by regulating the rate of rRNA synthesis, we next asked how this regulation was achieved? Previous studies had correlated LncRNA and pRNAs synthesis with CpG methylation and rDNA silencing (Mayer et al., 2006; Savic et al., 2014). However, when the rDNA methylation status was determined by WGB-Seq/EM-Seq or by inhibition of meCpG sensitive restriction enzyme (SmaI) cleavage we detected no significant change on TTF1 depletion in either conditional cell line (Figure 4B to D and S6A and B). In particular, neither the levels of methylation of the key regulatory CpG at −133 bp in the 47S Promoter (Santoro and Grummt, 2001) nor of 16 SmaI sites across the 18S and 28S genes were affected by TTF1 depletion. Direct determination of rDNA silencing by Psoralen Accessibility Crosslinking (PAC) (Conconi et al., 1989; Moss et al., 2019) also showed no change in the numbers of active rDNA copies on TTF1 depletion (Figure 4F and G). Further, the rDNA copy number determined both by WGS and by Southern blot also did not vary with TTF1 depletion (Figure 4E and S6B to D). Thus, the suppression of rRNA synthesis induced by TTF1 depletion were neither mediated by rDNA silencing, whether methylation dependent or not, nor by a reduction in rDNA copy number.

### LncRNA transcription complexes traverse the 47S Promoter

Since, neither rDNA silencing nor changes in gene copy number explained the downregulation of rRNA synthesis engendered by TTF1 depletion, we used DChIP-Seq to determine its effects on the engagement of the RPI across the rDNA. Depletion of TTF1 in both the C8 and C14 cells significantly reduced its recruitment at the Tsp, T_0_ and T_1-10_ rDNA binding sites, and this corresponded with a strong reduction in RPI associated with the Spacer Promoter and with increased RPI mapping throughout the adjacent enhancer repeats, as well as well downstream of the 47S pre-rRNA termination site (Figures 5A and B, and S7A and B). Thus, the RPI mapping data were fully consistent with the detection of LncRNA and pRNAs under these conditions (Figure 2B and S3) and confirmed that TTF1 was responsible for arresting transcription from the Spacer Promoter. The RPI mapping data further showed that on TTF1 depletion, the release of Spacer Promoter transcription complexes allowed their elongation well into the 47S Promoter (e.g. shading Figure 5B), and this was consistent with the detection of pRNA transcripts crossing the 47S Promoter (Prom2 amplicon in Figures 2B and S3). S1 RNA protection assays provided definitive evidence these pRNA transcripts were contiguous with the LncRNA derived from the Spacer Promoter since they traversed not only the 47S Promoter proximal T_0_ termination site but also continued right across the 47S Promoter and well into the downstream pre-rRNA transcribed region (Figure 5C and S8A to C).

**Figure 5.**
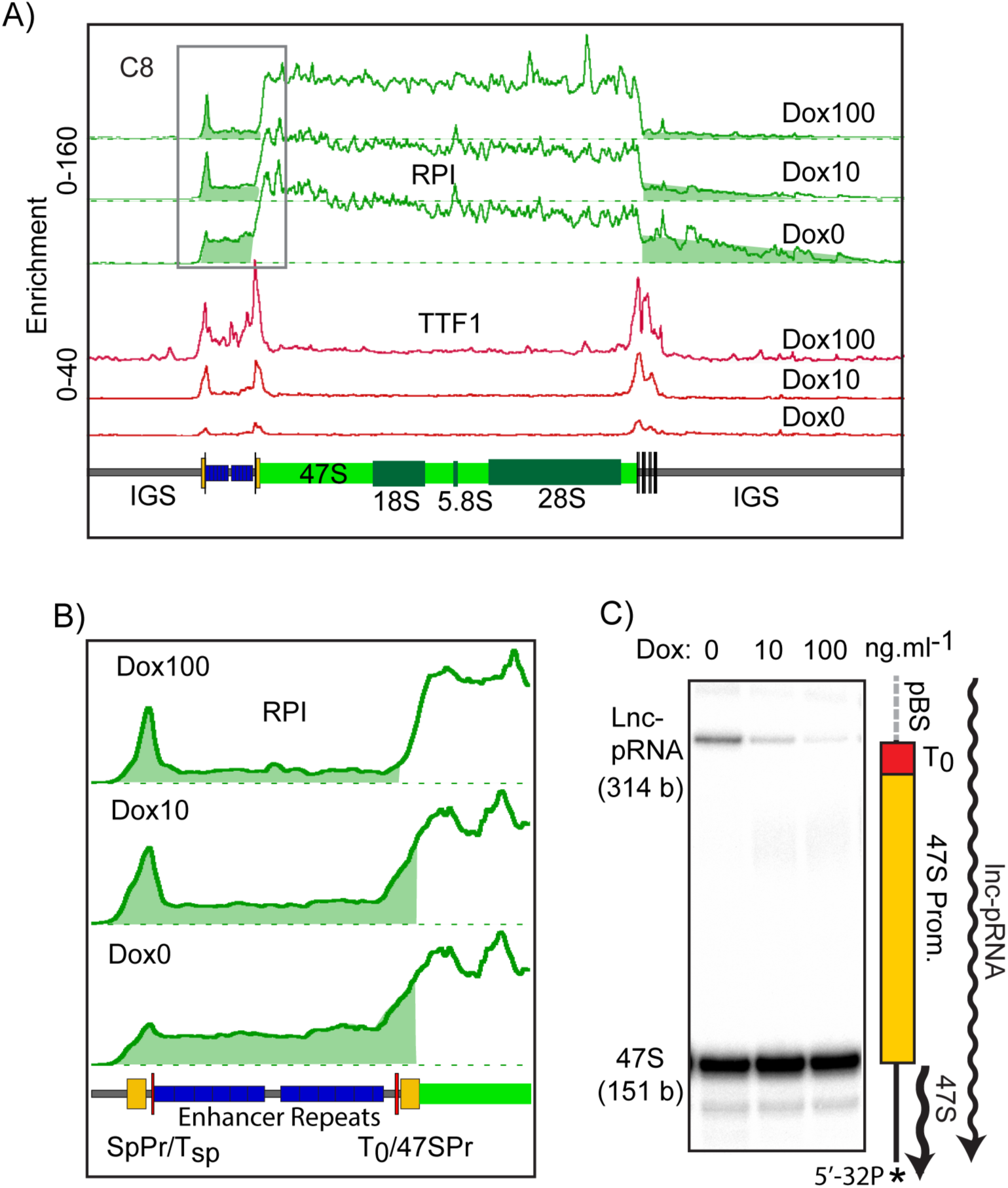
LncRNA elongation complexes readthrough the 47S Promoter in the absence of TTF1. A) RPI and TTF1 DChIP-Seq occupancy profiles across the rDNA repeat of clone C8 maintained at minimal (Dox0), suboptimal (Dox10) or optimal (Dox 100) TTF1 expression levels. B) Expanded view of the promoter and enhancer regions in A). In A and B reduced RPI occupancy at the Spacer Promoter and enhanced occupancy over the enhancer repeats and 47S Promoter, as well as readthrough of T_1-10_ site are indicated by shading. For an equivalent DChIP-Seq analysis of clone C14 see Figure S7. (C) S1 nuclease RNA protection assays detected a strong enhancement of LncRNA/pRNA transcripts passing through the 47S Promoter and into 47S pre-rRNA coding sequence of the rDNA on TTF1 depletion in clone C8, (see Figure S8 for more detail and quantification).

### Depletion of TTF1 limits RPI PIC formation at the 47S promoter

We argued that penetration of the Spacer Promoter transcripts into the functional 47S Promoter could interfere with its ability to efficiently promote pre-rRNA synthesis and might explain the suppression of rRNA synthesis on TTF1 depletion. To determine if this were the case, we determined 47S pre-initiation complex (PIC) formation before and after TTF1 depletion. Quantitative ChIP-qPCR and DChIP-Seq mapping assays revealed a reduction in SL1 (TAF1B) and UBTF occupancy at the 47S Promoter, consistent with the cooperative recruitment of these factors (Figure 6A and B) (Tremblay et al., 2022). We also noted a reduction in the Spacer Promoter PIC that was most probably related to the drastic loss of local RPI complexes arrested before promoter release (e.g. see Figure S9). Importantly, the reduction in 47S PIC formation corresponded closely with the reduced 47S pre-rRNA synthesis under the same conditions, both PIC levels and synthesis being suppressed by ∼50% (compare Figure 3E and TAF1B/UBTF promoter binding in Figure 6A and B). These data argued that TTF1 controls rRNA synthesis by regulating formation or stability of Preinitiation Complexes at the 47S Promoter and suggested a scenario in which transcription from the Spacer Promoter interfered with or occluded the function of this promoter (Figure 6C), see the Discussion.

**Figure 6.**
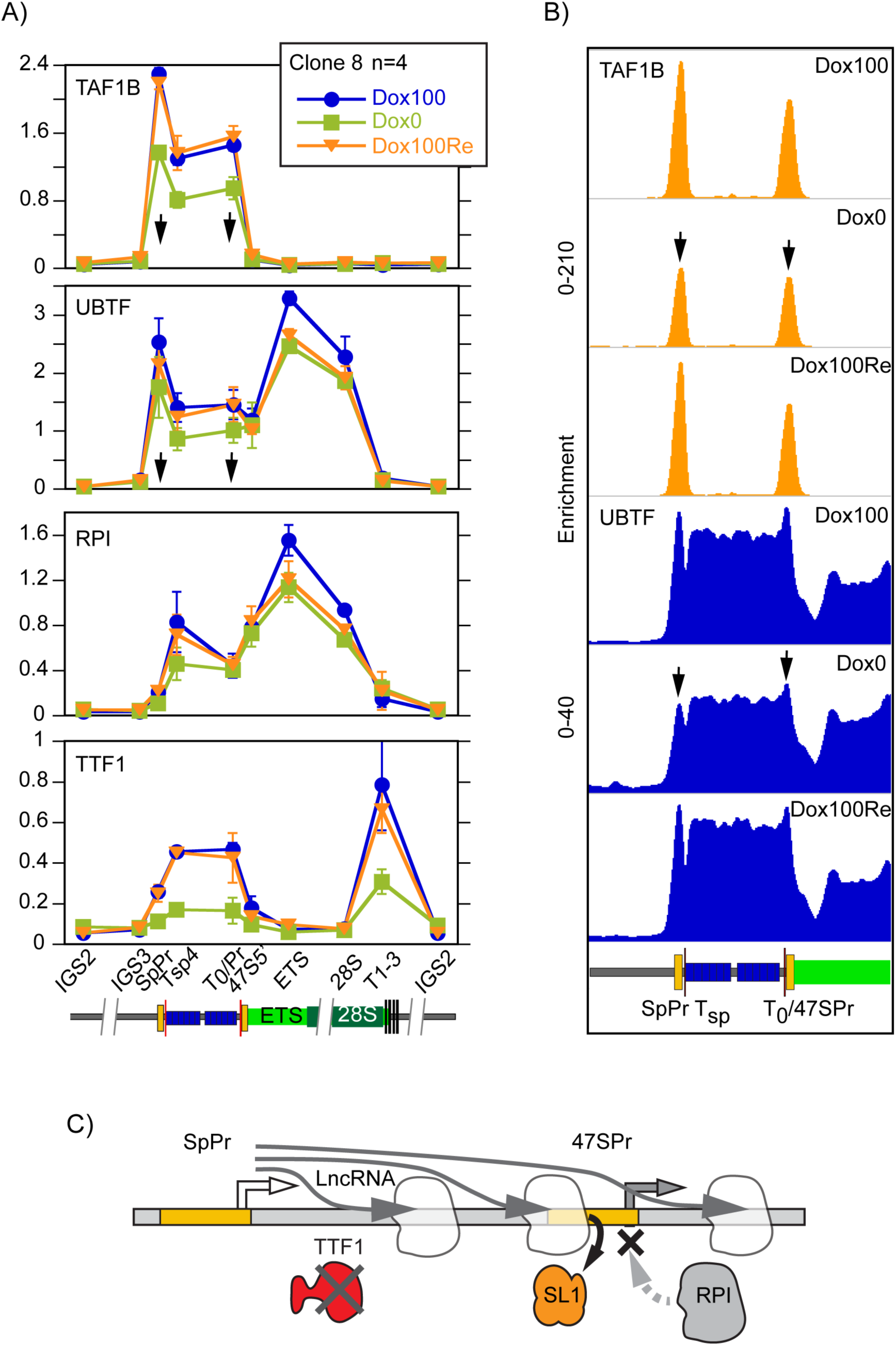
TTF1 depletion suppresses RPI PIC formation at the 47S Promoter. A) ChIP-qPCR and DChIP-Seq analysis of RPI, UBTF, TAF1B (SL1) and TTF1 occupancy across the rDNA in clone C8 either optimally expressing TTF1 (Dox100), TTF1 depleted for 7 days (Dox0) or 7 days after TTF1 reinduction (Dox100Re). The arrows indicate reduced recruitment of TAF1B and UBTF under TTF depletion (Dox0). The amplicons used and their positions across the rDNA are indicated below the mapping data. “n” Indicates the number of biological replicas and error bars the SEM. B) DChIP mapping of TAF1B and UBTF under the conditions used in A). For clarity only the rDNA region encompassing the Spacer and 47S Promoters is displayed. The full width DChIP mapping and including RPI and TTF1 can be found in Figure S9. C) Model of 47S Promoter interference/occlusion consistent with the TTF1 depletion data, showing LncRNA synthesis leading to the partial displacement of SL1 from the 47S Promoter and hence the suppression of RPI recruitment and 47S rRNA synthesis.

### p19ARF accumulation enhances LncRNA expression and RNA depletion in late passage MEFs

We had previously shown that nucleolar localization and rDNA binding by TTF1 was regulated by the p14/p19ARF (ARF) tumor suppressor, resulting in reduced pre-rRNA synthesis (Lessard et al., 2010). We now show that TTF1 levels also regulate cell growth and the cellular ribosome complement by determining the level of pre-rRNA synthesis. The combined data then suggested that the ARF tumour suppressor regulates cell growth via this same pathway. We argued that if this were the case, the natural accumulation of ARF and the reduced cellular proliferation occurring during senescence induced by passaging of Mouse Embryonic Fibroblasts (pMEFs) would necessarily induce LncRNA synthesis and would correlate with ribosome depletion. To test this possibility, we followed LncRNA and pRNA synthesis during the passaging of pMEFs. ARF accumulated in the pMEFs from passage 3 on, peaking from passage 4 to 6 before reducing somewhat at passage 7, by which time the cells were no longer detectably proliferating (Figure 7A). The corresponding levels of the LncRNA, pRNA and T_1-10_ readthrough were all observed to increase from passage 5 on and were 5- to 6-fold enhanced by passages 6 and 7 (Figure 7B). Strikingly, total RNA and hence ribosome levels were also reduced by 70 to 80% in late passage pMEFs (Figures 7C). Both this and the reduced proliferation of the late passage pMEFs paralled the suppression of ribosome biogenesis and proliferation observed on TTF1 depletion (Figure 3A and B). Hence, tumour suppression by ARF is fully consistent with its ability to displace TTF1 from the rDNA.

**Figure 7.**
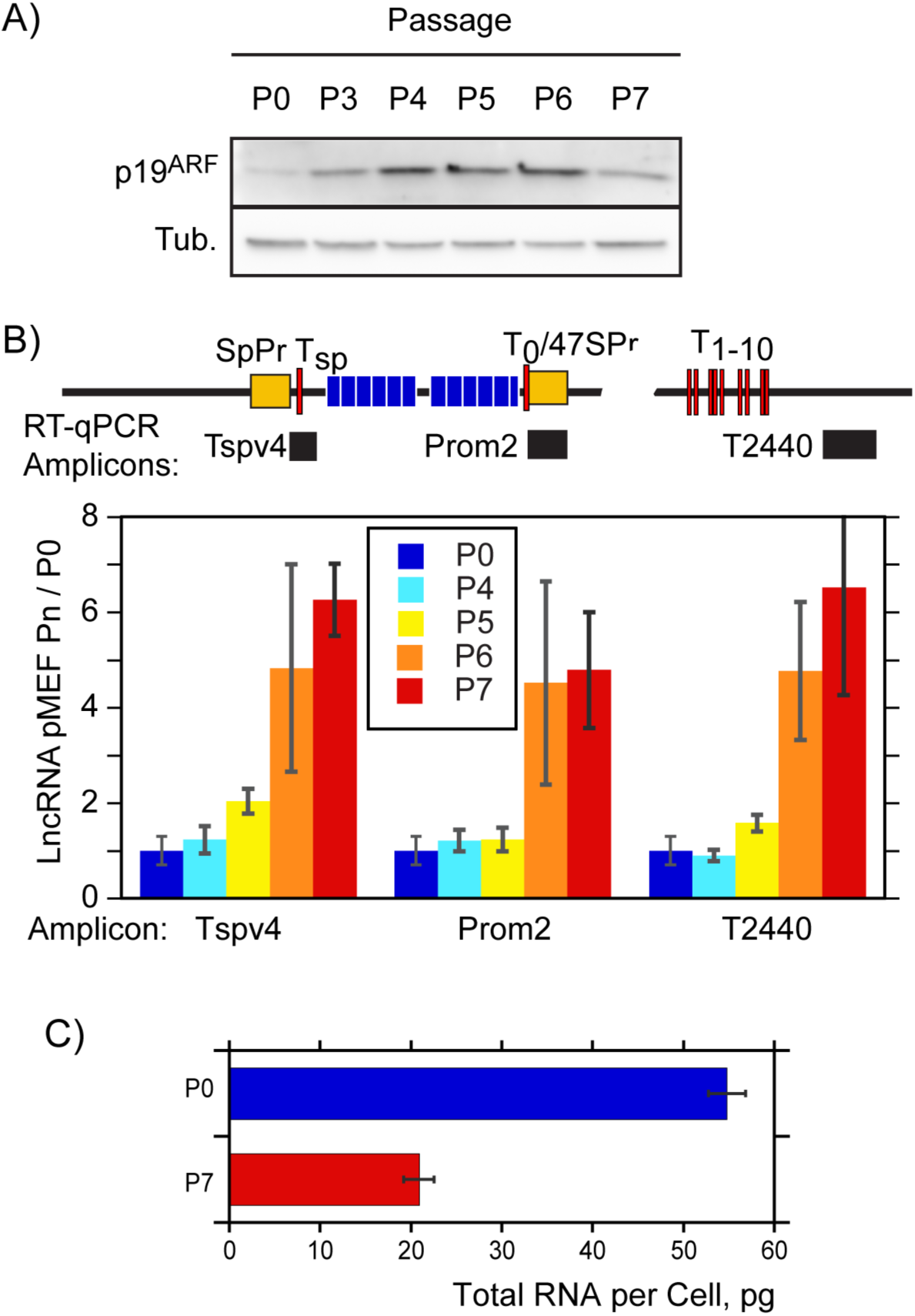
Enhanced LncRNA expression and strong depletion of total cellular RNA occurs concomitant with p19ARF accumulation in late passage MEFs. A) Western blot analysis of p19^ARF^ accumulation during replicative senescence, and B) RT-qPCR analysis of LncRNA and pRNA expression at sequential passages of wild type primary MEFs. “Tub.” refers to tubulin used as loading control in A), P0 to P7 refer to the culture passages analyzed, and the RT-qPCR amplicons used in B are indicated as in Figure 2B. C) Mean quantity of total RNA per primary MEF at passages 0 (P0) and 7 (P7). “n” Indicates the number of biological replicas and error bars the SEM.

## DISCUSSION

By generating cell lines conditional for the RPI termination factor TTF1 we have revealed an unexpected function of this factor in controlling the synthesis of non-coding RNAs from the ribosomal RNA genes and in determining the cellular proliferation rate by regulating ribosome biogenesis. The data further delineate a novel pathway of p53-independent tumour suppression downstream of p19ARF. Earlier data identified this tumour suppressor as a regulator of rRNA synthesis and of TTF1 nucleolar transport (Lessard et al., 2010; Sugimoto et al., 2003). We now show that in fact the nucleolar function of TTF1 is not limited to the termination of the primary rRNA transcript but also controls the synthesis of a regulatory LncRNA, and in so doing determines ribosome biogenesis and cell growth. Our data show that this LncRNA is in fact generated from the upstream Spacer Promoter duplications present in the ribosomal RNA genes (rDNA) of mouse but that are also found in human and indeed in most higher-eukaryote rDNAs (Moss et al., 2007). By binding at a site immediately downstream of the mouse Spacer Promoter and conserved from rodents to human, TTF1 arrests transcription from this promoter. This arrest occurs 30 or so nucleotides downstream of the Spacer Promoter and leads to a characteristic accumulation of RPI (Mars et al., 2018). Conditional depletion of TTF1 releases this transcriptional arrest and allows the Spacer Promoter to generate the LncRNA observed previously (Mayer et al., 2006; Santoro et al., 2010; Savic et al., 2014). Strikingly, synthesis of this non-coding RNA from its natural promoter corresponds to a significant slowing of the cellular division rate and the strong depletion of total cellular RNA and hence of ribosomes. These effects are explained by a reduction in the *de novo* synthesis of the 47S pre-rRNA and are fully reversed by the re-expression of TTF1. Together, our observations are consistent with a direct relationship between the cellular ribosome complement and the cellular proliferation rate as described by the cellular “growth laws” (Dai and Zhu, 2020; Scott et al., 2014), and may represent the first demonstration that this relationship is also valid in mammalian cells. Further, our data are consistent with regulation of TTF1 by p19ARF and hence delineate a novel and direct pathway between this tumour suppressor, ribosome biogenesis and the suppression of cell growth.

Previous studies had implicated TTF1 in rDNA silencing and shown that ectopic expression of the LncRNA and the pRNA can drive rDNA methylation (Mayer et al., 2006; Savic et al., 2014). In contrast, we find the “natural” *in cis* generation of these non-coding RNAs from the Spacer Promoter neither drives rDNA methylation nor silencing, excluding these explanations of the effects of TTF1 depletion. Further, this depletion does not affect rDNA copy number, nor does it affect genome-wide gene expression, strongly suggesting that TTF1 regulation of rRNA synthesis occurs via direct action with the rDNA itself. In fact, we find the reduction in *de novo* rRNA synthesis induced by TTF1 depletion is quantitatively explained by a reduction in RPI Preinitiation Complex (PIC) formation. Several possible explanations for this inhibition/destabilization are suggested by previous studies. However, the induction of gene silencing by DNA methylation, is clearly excluded by our analyses of both these parameters (Figures 4 and S6). TTF1 binding at the 47S Promoter has also been shown to displace nucleosomes and enhance promoter access (Längst et al., 1998), but as we have shown there is little evidence for nucleosomes or histone modifications on the rDNA promoters (Herdman et al., 2017; Moss et al., 2019). We cannot formally exclude other mechanisms such as TTF1 stabilization of the RPI PIC by direct contacts at the 47S Promoter, such interactions are absent from available TTF1 interactomes (BioGRID and IntAct (Del Toro et al., 2022; Oughtred et al., 2021)). However, the fact that TTF1 depletion allows Spacer Promoter transcripts to extend into the 47S Promoter strongly suggests that promoter interference or occlusion is likely to play a role (Adhya and Gottesman, 1982; Shearwin et al., 2005; Zhang and Bremer, 1996). In this scenario RPI transcription from the Spacer Promoter, by reading into and through the downstream 47S promoter, would displaces or destabilize the UBTF/SL1 PIC and hence reduce the rate of productive rRNA synthesis (Figure 6C). As such, in this scenario it would be the act of LncRNA transcription rather than the generation of the LncRNA itself that would be the key factor.

TTF1 control of 47S Promoter occlusion is also fully consistent with the known actions of the yeast ortholog Reb1, which performs an important failsafe ‘roadblock” termination function to prevent readthrough interference between adjacent yeast genes (Candelli et al., 2018; Colin et al., 2014). In fact, the potential for transcriptional interference in the context of rDNA promoter duplications was recognized many years ago (Firek et al., 1989; Grummt et al., 1986; McStay and Reeder, 1990; Mitchelson and Moss, 1987; Moss, 1983; Moss et al., 1992). However, its regulatory importance was at the time unclear. Our data now provide a *raison d’être* for the existence of spacer promoter duplications by showing that, in combination with TTF1 roadblock termination, they can control the rate of ribosome biogenesis and hence the rate of cell proliferation. This mechanism might also have broader implications in the transitions between active and inactive rDNA copies that must inevitably occur during development and differentiation.

## ACKNOWLEDEGEMENTS

This work was supported by an operating grant from the Canadian Institutes of Health Research (CIHR), [grant number MOP12205/PJT153266]. The Research Centre of the Québec University Hospital Centre (CRCHU de Québec-Université Laval) is supported by the Fonds de Recherche du Québec - Santé (FRQS). We also thank Dr Andre Nussenzweig for his aid with early ChIP-Seq analyses and Dr Steve Bilodeau (Cancer Division, CRCHU de Québec-Université Laval) for his help with RNA-Seq analyses.

