## Supplemental Figures for "TTF1 control of LncRNA synthesis and cell growth delineates a tumour suppressor pathway acting directly on the ribosomal RNA Genes"

### SUPPLEMENTAL FIGURE LEGENDS

**Figure S1. Generation of conditional mouse TTF1 cell lines under doxycycline control.** A) design of the CRISPR/Cas9 guide RNA to target exon 2 of endogenous TTF1 and the silent mutation introduced into the 3xFLAG-TTF1 cDNA to render it resistant to deletion. B) Western analysis of doxycycline (Dox) induced 3xFLAG-TTF1 expression in clones C8 and C14 revealed with the anti-TTF1 and anti-FLAG antibodies. C) Mean steady-state 3xFLAG-TTF1 expression levels in C8 and C14 after Dox induction and in NIH3T3 estimated from Western blot analyses using the anti-TTF1 antibody. The data derive from 8 biological replicas and error bars show the SEM. D) ChIP-qPCR analysis of TTF1 occupancy at sites across the rDNA in C8 and C14 either with Dox induction or 3 days after Dox withdrawal. The amplicons used and their positions across the rDNA are indicated below the mapping data.

**Figure S2. Sequences of the wild type and CRISPR/Cas9 deleted TTF1 alleles.** The sequences across Exon 2 of the wild type (wt) Ttf1 gene and the deleted alleles in clones C8 and C14.

**Figure S3. LncRNA expression in clone c14.** Upper panel; organisation of the mouse rDNA locus indicating the positions of the Spacer (SpPr) and 47S (47S Pr) promoter sequences and the canonical TTF1 binding sites Tsp, $T_0$ and $T_{1\text{-}10}$. The positions of the RT-qPCR amplicons are indicated as in Figure 2B. Lower panel; RT-qPCR analysis of RNA from clone C14 grown under the indicated conditions of TTF1 induction (Dox0 and 10 refer respectively to growth with 0 or $10ng.ml^{-1}$ Dox). Data were normalized to RPS12 mRNA levels (Savic et al., 2014) and are presented relative to wild type NIH3T3. The data derive from 4 independent biological replicas (n = 4) each analyzed in triplicate, and error bars indicate the SEM.

**Figure S4. Dox induction of TTF1 determines cell proliferation and 47S pre-rRNA synthesis rates, and the cellular ribosome complement.** A) and B) Growth curves for the C8, C14 and NIH3T3 under standard condition with or without Dox induction of the TTF1 transgene. The doubling times indicated were estimated from the $\log_2$ plots by linear curve fit, "n" refers to the number of biological replicates and error bars indicate the SEM. C) Summary of doubling times for the C14 cell line taken from B. D) Example of the reversibility of TTF1 depletion on cell proliferation. The C8 clone was grown in 100 $ng.ml^{-1}$ doxycycline (+Dox) for 6 days. Doxycycline was then removed (-Dox) and cells maintained in culture until day 10 at which time 100 $ng.ml^{-1}$ doxycycline (+ Dox) was re-added and cells maintained in culture for another 5 days. E) Cellular RNA content of NIH3T3 cells and C14 cells grown in the absence or presence of 10 $ng.ml^{-1}$ doxycycline. F) 47S pre-rRNA synthesis rates in the C14 cell line grown in the absence or presence of 10 $ng.ml^{-1}$ doxycycline relative to NIH3T3. The data were normalized to the steady state bulk 28S rRNA (left panel) or to the cellular RNA content (right panel). "n" Indicates the number of biological replicas and error bars the SEM.

**Figure S5. TTF1 depletion does not significantly affect genome-wide gene expression.** A) Western blot analysis of TTF1 levels in clone C8 maintained in 100$ng.ml^{-1}$ doxycycline (Dox100 Day0), or grown for 3, 5 or 7 days without doxycycline (Dox0) before doxycycline readdition for a further 3, 5 or 7 days (Dox100 readd). RNAs from cultures analyzed in A were then subjected to RNA-Seq analysis to determine genome-wide changes in gene expression. B) and C) Mean Difference Plots of mRNA expression against mean expression level (CPM) for biological duplicate RNA samples analyzed in Figure 4A. No significant changes in gene expression were detected due to TTF1 depletion or its reestablishment. This is illustrated by the results for the RPI transcription factor genes and for the ribosomal protein genes Rps5 and Rps12 used in qPCR analyses of lncRNA expression.

**Figure S6. Determination of CpG methylation and rDNA copy number before and after TTF1 depletion.** A) Typical examples of WGB-Seq/EM-Seq mapping showing the percentage of meCpG across the rDNA repeat in NIH3T3 and clone C8 depleted of TTF1 or optimally expressing TTF1 (Dox0, Dox100). The upper panel shows mapping across the full rDNA repeat and the Lower panel detailed mapping of the 47S Promoter and immediate flanking regions. B) Upper panel shows an example of Southern blotting of a 4.8kbp BamHI fragment spanning the 18S and 28S rDNA coding region subjected to SmaI or XmaI cleavage to detect methylation levels at 5'CCCGGG sequences. The lower panel indicates the positions of cleavage sites within the 4.8kbp BamHI fragment, the PflMI-BamHI probe and the 7.9 kbp frament from the UBTF gene used as single copy reference. C) and D) Respectively the rDNA copy numbers estimated from Whole Genome Sequencing (WGS) and from Southern blotting. "n" indicates the number of biological replicas and error bars the SEM.

**Figure S7. LncRNA elongation complexes also readthrough to the 47S Promoter in clone C14 on TTF1 depletion.** A) RPI and TTF1 DChIP-Seq occupancy profiles across the rDNA repeat of clone C14 maintained at minimal (Dox0) or optimal (Dox 10) TTF1 expression levels. B) Expanded view of the promoter and enhancer regions in A). In A and B reduced RPI occupancy at the Spacer Promoter and enhanced occupancy over the enhancer repeats and 47S Promoter, as well as readthrough of $T_{1\text{-}10}$ site are indicated by shading.

**Figure S8. S1 protection mapping reveal LncRNA transcripts traversing the 47S Promoter.** A) S1 nuclease RNA protection assays of RNA from clone C8 detected strong enhancement of LncRNA transcripts when grown without doxycycline (Dox0) as compared to growth in suboptimal or optimal doxycycline (Dox10 or 100). B) Organization of the S1 probe showing the 47S pre-RNA, the presumed LncRNA/pRNA and construction of the probe that includes a rDNA non-complementary 3' region (pBS) and 5’ labelling site. The LncRNA/pRNA transcripts protect the full length of the rDNA complementary probe region from the upstream boundary of the 47S Promoter to 151 b. into 47S pre-rRNA coding region. C) Quantitation of probe protection by the LncRNA/pRNA transcripts normalized to the signal for 47S pre-rRNA protection. In A and C "Days" refer to the culture time in the absence of doxycycline, "n" indicates the number of biological replicas and error bars the SEM.

**Figure S9. DChIP mapping of RPI, TAF1B, UBTF and TTF1 before, after TTF1 depletion.** The figure shows the occupancy of the four factors across the full width of the rDNA in clone C8 corresponding to the data shown in Figure 6B. C8 cells were maintained for >7 days with TTF1 induction (Dox100), 7 days with no TTF induction (Dox0) and finally 7 days after re-induction of TTF1 (Dox100Re).

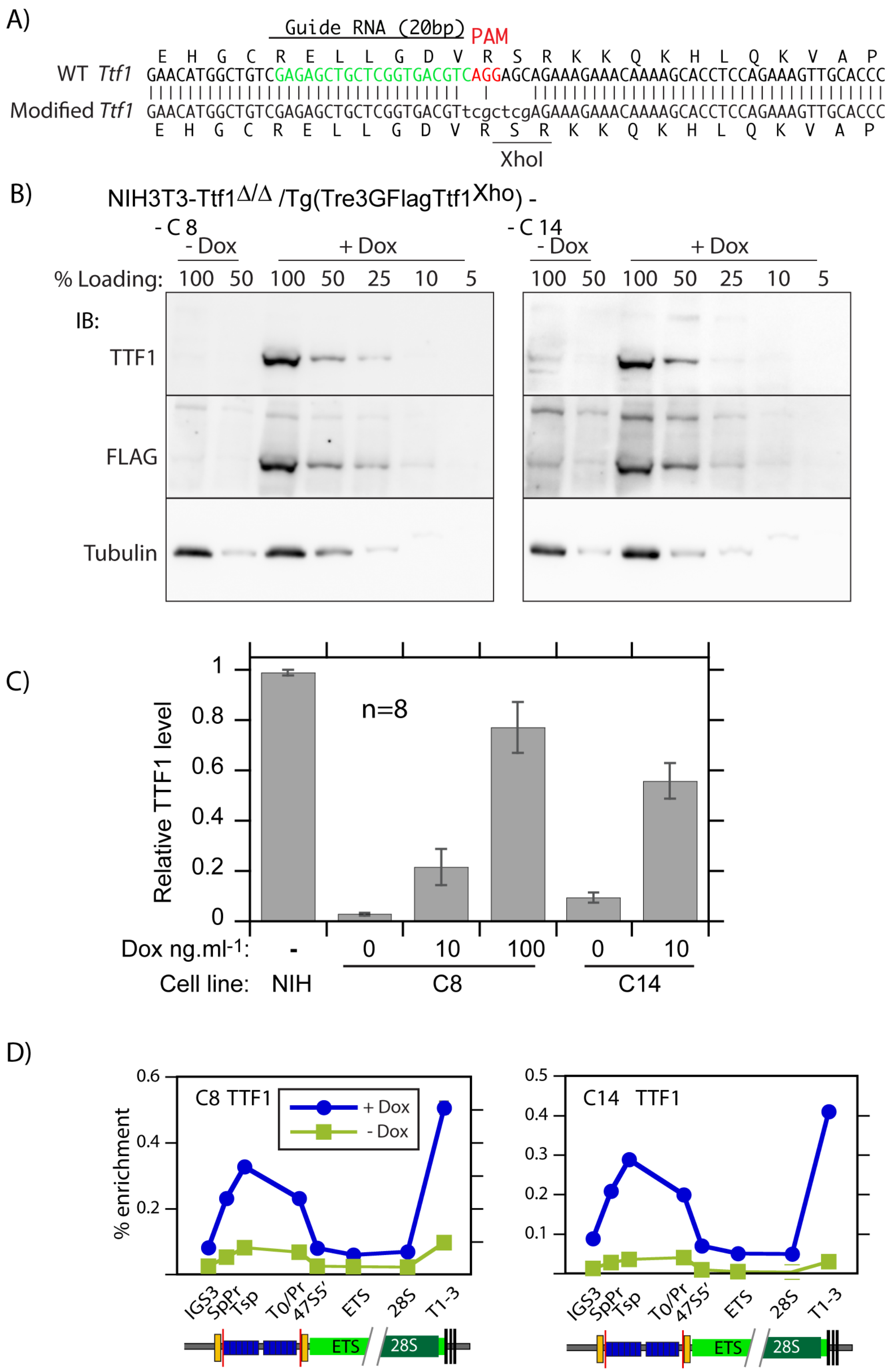
A)
Guide RNA (20bp)
PAM
E H G C R E L L G D V R S R K K Q K H L Q K V A P
WT Ttf1 GAACATGGCTGTCGAGAGCTGCTCGGTGACGTCAGGAGCAGAAAGAAACAAAAGCACCTCCAGAAAGTTGCACCC
Modified Ttf1 GAACATGGCTGTCGAGAGCTGCTCGGTGACGTtcgctcgAGAAAGAAACAAAAGCACCTCCAGAAAGTTGCACCC
E H G C R E L L G D V R S R K K Q K H L Q K V A P
XhoI
B)
NIH3T3-Ttf1Δ/Δ /Tg(Tre3GFlagTtf1Xho) -
- C 8
- C 14
- Dox
+ Dox
% Loading: 100 50 100 50 25 10 5
IB:
TTF1
FLAG
Tubulin
C)
Relative TTF1 level
n=8
1
0.8
0.6
0.4
0.2
0
Dox ng.ml-1: - 0 10 100 0 10
Cell line: NIH C8 C14
D)
% enrichment
C8 TTF1
C14 TTF1
+ Dox
- Dox
0.6
0.4
0.2
0.5
0.4
0.3
0.2
0.1
IGS3
SpPr
Tsp
To/Pr
47S5'
ETS
28S
T1-3
ETS
28S

Figure S1

### Endogenous TTF wt alleles

Exon 2 ATG

GGACAGGATGAAAGGGGGCACAAGCAAATTTAAGACCCACACTGAAACTTTGTACAAGAAAAAGAAATGGTCTTCTGTGTCTGAAAAAAGACCTCAGAAATGTCCCTCTCAGTGCTTGGAGAGCAAGCAGCCTCAGGTATCTGTGTTGGGGAAGAGGAGGAG

▶ M K G G T S K F K T H T E T L Y K K K K W S S V S E K R P Q K C P S Q C L E S K Q P Q V S V L G K R R R

GGCGTCCCAGACACCAGCCCAGGAGACCTTAGAGAGTGAGTGGCCTCAAAAGGCCAAGAGGAAGAAGAGGAGGAGGGAACCCCAGACACCAGCCCAGGAGACCTTAGAGAGTGAGTGGCCTCAAAAGGCCAAGAAGAAGAAGAGAAGGGGGGAACCCCAGAC

▶ A S Q T P A Q E T L E S E W P Q K A K R K K R R R E P Q T P A Q E T L E S E W P Q K A K K K K R R G E P Q T

ACCAACCCAAGAGAGTTTGGAGAGTGAGCAGCCTCCAGTGTCCTTGCTGGGGAAGAGAAGGAGGGAATCCCAGACACCAGCCCAGGAGAACTCGGAGAGTGAGCAGCCTCGGAAAGCCAAGAGAAGGAGGAAGAAAAGGAAAGGGTCCCAGCAACCAACTTC

▶ P T Q E S L E S E Q P P V S L L G K R R R E S Q T P A Q E N S E S E Q P R K A K R R R K K R K G S Q Q P T S

CTCCCTTCTGAAAACTCCAGAAACCTTTCTTAAAGCTAAGAAAACCACTTCTGCCCACAAGAAGAAAAAGAATAGTGTTCTGGAGGTGGATATGGAAACTGGGATCATCCTTGTAGATAAAGAAAACATGGAGAACCTGCTAGAGACTTCTAGAAAGGATGT

▶ S L L K T P E T F L K A K K T T S A H K K K K N S V L E V D M E T G I I L V D K E N M E N L L E T S R K D V

GGATATTGTTTATGTTGATATGAGCAAGGGACAAAGGTCAGCAAAAGTGCGTGAAACAGGAGAGCTGCCCGCTGCTAAGCCACAGGAACATGGCTGTCGAGAGCTGCTCGGTGACGTCAGGAGCAGAAAGAAACAAAAGCACCTCCAGAAAGTTGCACCCTG

▶ D I V Y V D M S K G Q R S A K V R E T G E L P A A K P Q E H G C R E L L G D V R S R K K Q K H L Q K V A P W

GGATGTTGTTCAGGGGAGCCAGCCTGAGAGTATCTCCCTGCCCCCATCAGAGCCCTTGTCTTCTGAAGATTTAGAAGGCAAAAGCACAGAAGCAGCAGTGTTTTGTAAAAAGAAGTCTAAAAAAAACGTGTTCAGGAGCCAGGAATTGGAGCCCATCCCCGA

▶ D V V Q G S Q P E S I S L P P S E P L S S E D L E G K S T E A A V F C K K K S K K N V F R S Q E L E P I P D

CAGTCTTGATGACTCGGAGACCATCTCTGAGAGACTTGATAGCACACACCATGGAGGAGCTGTCGGGGCTGGAGAAGAATGTGAGAGCACCAAAGAATCCCACAGTATCAAAAAAAAGTCCAAGAAAAAGAAGCACAAGTCTGTTGCTCTTGCCACATCTAG

▶ S L D D S E T I S E R L D S T H H G G A V G A G E E C E S T K E S H S I K K K S K K K K H K S V A L A T S S

TGACAGTGCCTCAGTGACAGACAGCAAGGCTAAGAATGCCCTGGTGGACTCCTCAGAAGGCAGTGGTGCTGTGAGGGAAGAGGACGTGGACCACAGGCCAGCAGAGGCAGAAGCCCAGGCTTGTTCCACTGAGAAGCACAGGGAAGCCATGCAGAG

▶ D S A S V T D S K A K N A L V D S S E G S G A V R E E D V D H R P A E A E A Q A C S T E K H R E A M Q

### Clone 8; Endogenous TTF allele 1 (+1bp)

Exon 2 ATG

GGACAGGATGAAAGGGGGCACAAGCAAATTTAAGACCCACACTGAAACTTTGTACAAGAAAAAGAAATGGTCTTCTGTGTCTGAAAAAAGACCTCAGAAATGTCCCTCTCAGTGCTTGGAGAGCAAGCAGCCTCAGGTATCTGTGTTGGGGAAGAGGAGGAG

▶ M K G G T S K F K T H T E T L Y K K K K W S S V S E K R P Q K C P S Q C L E S K Q P Q V S V L G K R R R

GGCGTCCCAGACACCAGCCCAGGAGACCTTAGAGAGTGAGTGGCCTCAAAAGGCCAAGAGGAAGAAGAGGAGGAGGGAACCCCAGACACCAGCCCAGGAGACCTTAGAGAGTGAGTGGCCTCAAAAGGCCAAGAAGAAGAAGAGAAGGGGGGAACCCCAGAC

▶ A S Q T P A Q E T L E S E W P Q K A K R K K R R R E P Q T P A Q E T L E S E W P Q K A K K K K R R G E P Q T

ACCAACCCAAGAGAGTTTGGAGAGTGAGCAGCCTCCAGTGTCCTTGCTGGGGAAGAGAAGGAGGGAATCCCAGACACCAGCCCAGGAGAACTCGGAGAGTGAGCAGCCTCGGAAAGCCAAGAGAAGGAGGAAGAAAAGGAAAGGGTCCCAGCAACCAACTTC

▶ P T Q E S L E S E Q P P V S L L G K R R R E S Q T P A Q E N S E S E Q P R K A K R R R K K R K G S Q Q P T S

CTCCCTTCTGAAAACTCCAGAAACCTTTCTTAAAGCTAAGAAAACCACTTCTGCCCACAAGAAGAAAAAGAATAGTGTTCTGGAGGTGGATATGGAAACTGGGATCATCCTTGTAGATAAAGAAAACATGGAGAACCTGCTAGAGACTTCTAGAAAGGATGT

▶ S L L K T P E T F L K A K K T T S A H K K K K N S V L E V D M E T G I I L V D K E N M E N L L E T S R K D V

GGATATTGTTTATGTTGATATGAGCAAGGGACAAAGGTCAGCAAAAGTGCGTGAAACAGGAGAGCTGCCCGCTGCTAAGCCACAGGAACATGGCTGTCGAGAGCTGCTCGGTGACCGTCAGGAGCAGAAAGAAACAAAAGCACCTCCAGAAAGTTGCACCCT

▶ D I V Y V D M S K G Q R S A K V R E T G E L P A A K P Q E H G C R E L L G D R Q E Q K E T K A P P E S C T L

GGGATGTTGTTCAGGGGAGCCAGCCTGAGAGTATCTCCCTGCCCCCATCAGAGCCCTTGTCTTCTGAAGATTTAGAAGGCAAAAGCACAGAAGCAGCAGTGTTTTGTAAAAAGAAGTCTAAAAAAAACGTGTTCAGGAGCCAGGAATTGGAGCCCATCCCCG

▶ G C C S G E P A

ACAGTCTTGATGACTCGGAGACCATCTCTGAGAGACTTGATAGCACACACCATGGAGGAGCTGTCGGGGCTGGAGAAGAATGTGAGAGCACCAAAGAATCCCACAGTATCAAAAAAAAGTCCAAGAAAAAGAAGCACAAGTCTGTTGCTCTTGCCACATCTA

GTGACAGTGCCTCAGTGACAGACAGCAAGGCTAAGAATGCCCTGGTGGACTCCTCAGAAGGCAGTGGTGCTGTGAGGGAAGAGGACGTGGACCACAGGCCAGCAGAGGCAGAAGCCCAGGCTTGTTCCACTGAGAAGCACAGGGAAGCCATGCAGAG

### Clone 8; Endogenous TTF allele 2 (-62bp)

Exon 2 ATG

GGACAGGATGAAAGGGGGCACAAGCAAATTTAAGACCCACACTGAAACTTTGTACAAGAAAAAGAAATGGTCTTCTGTGTCTGAAAAAAGACCTCAGAAATGTCCCTCTCAGTGCTTGGAGAGCAAGCAGCCTCAGGTATCTGTGTTGGGGAAGAGGAGGAG

▶ M K G G T S K F K T H T E T L Y K K K K W S S V S E K R P Q K C P S Q C L E S K Q P Q V S V L G K R R R

GGCGTCCCAGACACCAGCCCAGGAGACCTTAGAGAGTGAGTGGCCTCAAAAGGCCAAGAGGAAGAAGAGGAGGAGGGAACCCCAGACACCAGCCCAGGAGACCTTAGAGAGTGAGTGGCCTCAAAAGGCCAAGAAGAAGAAGAGAAGGGGGGAACCCCAGAC

▶ A S Q T P A Q E T L E S E W P Q K A K R K K R R R E P Q T P A Q E T L E S E W P Q K A K K K K R R G E P Q T

ACCAACCCAAGAGAGTTTGGAGAGTGAGCAGCCTCCAGTGTCCTTGCTGGGGAAGAGAAGGAGGGAATCCCAGACACCAGCCCAGGAGAACTCGGAGAGTGAGCAGCCTCGGAAAGCCAAGAGAAGGAGGAAGAAAAGGAAAGGGTCCCAGCAACCAACTTC

▶ P T Q E S L E S E Q P P V S L L G K R R R E S Q T P A Q E N S E S E Q P R K A K R R R K K R K G S Q Q P T S

CTCCCTTCTGAAAACTCCAGAAACCTTTCTTAAAGCTAAGAAAACCACTTCTGCCCACAAGAAGAAAAAGAATAGTGTTCTGGAGGTGGATATGGAAACTGGGATCATCCTTGTAGATAAAGAAAACATGGAGAACCTGCTAGAGACTTCTAGAAAGGATGT

▶ S L L K T P E T F L K A K K T T S A H K K K K N S V L E V D M E T G I I L V D K E N M E N L L E T S R K D V

GGATATTGTTTATGTTGATATGAGCAAGGGACAAAGGTCAGCAAAAGTGCGTGAAACAGGAGAGCTGCCCGCTGCTAAGCCACAGGAACATGGCACCCTGGGATGTTGTTCAGGGGAGCCAGCCTGAGAGTATCTCCCTGCCCCCATCAGAGCCCTTGTCTT

▶ D I V Y V D M S K G Q R S A K V R E T G E L P A A K P Q E H G T L G C C S G E P A

CTGAAGATTTAGAAGGCAAAAGCACAGAAGCAGCAGTGTTTTGTAAAAAGAAGTCTAAAAAAAACGTGTTCAGGAGCCAGGAATTGGAGCCCATCCCCGACAGTCTTGATGACTCGGAGACCATCTCTGAGAGACTTGATAGCACACACCATGGAGGAGCTG

TCGGGGCTGGAGAAGAATGTGAGAGCACCAAAGAATCCCACAGTATCAAAAAAAAGTCCAAGAAAAAGAAGCACAAGTCTGTTGCTCTTGCCACATCTAGTGACAGTGCCTCAGTGACAGACAGCAAGGCTAAGAATGCCCTGGTGGACTCCTCAGAAGGCA

GTGGTGCTGTGAGGGAAGAGGACGTGGACCACAGGCCAGCAGAGGCAGAAGCCCAGGCTTGTTCCACTGAGAAGCACAGGGAAGCCATGCAGAG

### Clone 14; Endogenous TTF allele 1 (-7bp)

Exon 2 ATG

GGACAGGATGAAAGGGGGCACAAGCAAATTTAAGACCCACACTGAAACTTTGTACAAGAAAAAGAAATGGTCTTCTGTGTCTGAAAAAAGACCTCAGAAATGTCCCTCTCAGTGCTTGGAGAGCAAGCAGCCTCAGGTATCTGTGTTGGGGAAGAGGAGGAG

▶ M K G G T S K F K T H T E T L Y K K K K W S S V S E K R P Q K C P S Q C L E S K Q P Q V S V L G K R R R

GGCGTCCCAGACACCAGCCCAGGAGACCTTAGAGAGTGAGTGGCCTCAAAAGGCCAAGAGGAAGAAGAGGAGGAGGGAACCCCAGACACCAGCCCAGGAGACCTTAGAGAGTGAGTGGCCTCAAAAGGCCAAGAAGAAGAAGAGAAGGGGGGAACCCCAGAC

▶ A S Q T P A Q E T L E S E W P Q K A K R K K R R R E P Q T P A Q E T L E S E W P Q K A K K K K R R G E P Q T

ACCAACCCAAGAGAGTTTGGAGAGTGAGCAGCCTCCAGTGTCCTTGCTGGGGAAGAGAAGGAGGGAATCCCAGACACCAGCCCAGGAGAACTCGGAGAGTGAGCAGCCTCGGAAAGCCAAGAGAAGGAGGAAGAAAAGGAAAGGGTCCCAGCAACCAACTTC

▶ P T Q E S L E S E Q P P V S L L G K R R R E S Q T P A Q E N S E S E Q P R K A K R R R K K R K G S Q Q P T S

CTCCCTTCTGAAAACTCCAGAAACCTTTCTTAAAGCTAAGAAAACCACTTCTGCCCACAAGAAGAAAAAGAATAGTGTTCTGGAGGTGGATATGGAAACTGGGATCATCCTTGTAGATAAAGAAAACATGGAGAACCTGCTAGAGACTTCTAGAAAGGATGT

▶ S L L K T P E T F L K A K K T T S A H K K K K N S V L E V D M E T G I I L V D K E N M E N L L E T S R K D V

GGATATTGTTTATGTTGATATGAGCAAGGGACAAAGGTCAGCAAAAGTGCGTGAAACAGGAGAGCTGCCCGCTGCTAAGCCACAGGAACATGGCTGTCGAGAGCTGCCGTCAGGAGCAGAAAGAAACAAAAGCACCTCCAGAAAGTTGCACCCTGGGATGTT

▶ D I V Y V D M S K G Q R S A K V R E T G E L P A A K P Q E H G C R E L P S G A E R N K S T S R K L H P G M L

GTTCAGGGGAGCCAGCCTGAGAGTATCTCCCTGCCCCCATCAGAGCCCTTGTCTTCTGAAGATTTAGAAGGCAAAAGCACAGAAGCAGCAGTGTTTTGTAAAAAGAAGTCTAAAAAAAACGTGTTCAGGAGCCAGGAATTGGAGCCCATCCCCGACAGTCTT

▶ F R G A S L R V S P C P H Q S P C L L K I

GATGACTCGGAGACCATCTCTGAGAGACTTGATAGCACACACCATGGAGGAGCTGTCGGGGCTGGAGAAGAATGTGAGAGCACCAAAGAATCCCACAGTATCAAAAAAAAGTCCAAGAAAAAGAAGCACAAGTCTGTTGCTCTTGCCACATCTAGTGACAGT

GCCTCAGTGACAGACAGCAAGGCTAAGAATGCCCTGGTGGACTCCTCAGAAGGCAGTGGTGCTGTGAGGGAAGAGGACGTGGACCACAGGCCAGCAGAGGCAGAAGCCCAGGCTTGTTCCACTGAGAAGCACAGGGAAGCCATGCAGAG

### Clone 14; Endogenous TTF allele 2 (-10bp)

Exon 2 ATG

GGACAGGATGAAAGGGGGCACAAGCAAATTTAAGACCCACACTGAAACTTTGTACAAGAAAAAGAAATGGTCTTCTGTGTCTGAAAAAAGACCTCAGAAATGTCCCTCTCAGTGCTTGGAGAGCAAGCAGCCTCAGGTATCTGTGTTGGGGAAGAGGAGGAG

▶ M K G G T S K F K T H T E T L Y K K K K W S S V S E K R P Q K C P S Q C L E S K Q P Q V S V L G K R R R

GGCGTCCCAGACACCAGCCCAGGAGACCTTAGAGAGTGAGTGGCCTCAAAAGGCCAAGAGGAAGAAGAGGAGGAGGGAACCCCAGACACCAGCCCAGGAGACCTTAGAGAGTGAGTGGCCTCAAAAGGCCAAGAAGAAGAAGAGAAGGGGGGAACCCCAGAC

▶ A S Q T P A Q E T L E S E W P Q K A K R K K R R R E P Q T P A Q E T L E S E W P Q K A K K K K R R G E P Q T

ACCAACCCAAGAGAGTTTGGAGAGTGAGCAGCCTCCAGTGTCCTTGCTGGGGAAGAGAAGGAGGGAATCCCAGACACCAGCCCAGGAGAACTCGGAGAGTGAGCAGCCTCGGAAAGCCAAGAGAAGGAGGAAGAAAAGGAAAGGGTCCCAGCAACCAACTTC

▶ P T Q E S L E S E Q P P V S L L G K R R R E S Q T P A Q E N S E S E Q P R K A K R R R K K R K G S Q Q P T S

CTCCCTTCTGAAAACTCCAGAAACCTTTCTTAAAGCTAAGAAAACCACTTCTGCCCACAAGAAGAAAAAGAATAGTGTTCTGGAGGTGGATATGGAAACTGGGATCATCCTTGTAGATAAAGAAAACATGGAGAACCTGCTAGAGACTTCTAGAAAGGATGT

▶ S L L K T P E T F L K A K K T T S A H K K K K N S V L E V D M E T G I I L V D K E N M E N L L E T S R K D V

GGATATTGTTTATGTTGATATGAGCAAGGGACAAAGGTCAGCAAAAGTGCGTGAAACAGGAGAGCTGCCCGCTGCTAAGCCACAGGAACATGGCTGTCGAGAGCTGCTCGGTGCAGAAAGAAACAAAAGCACCTCCAGAAAGTTGCACCCTGGGATGTTGTT

▶ D I V Y V D M S K G Q R S A K V R E T G E L P A A K P Q E H G C R E L L G A E R N K S T S R K L H P G M L F

CAGGGGAGCCAGCCTGAGAGTATCTCCCTGCCCCCATCAGAGCCCTTGTCTTCTGAAGATTTAGAAGGCAAAAGCACAGAAGCAGCAGTGTTTTGTAAAAAGAAGTCTAAAAAAAACGTGTTCAGGAGCCAGGAATTGGAGCCCATCCCCGACAGTCTTGAT

▶ R G A S L R V S P C P H Q S P C L L K I

GACTCGGAGACCATCTCTGAGAGACTTGATAGCACACACCATGGAGGAGCTGTCGGGGCTGGAGAAGAATGTGAGAGCACCAAAGAATCCCACAGTATCAAAAAAAAGTCCAAGAAAAAGAAGCACAAGTCTGTTGCTCTTGCCACATCTAGTGACAGTGCC

TCAGTGACAGACAGCAAGGCTAAGAATGCCCTGGTGGACTCCTCAGAAGGCAGTGGTGCTGTGAGGGAAGAGGACGTGGACCACAGGCCAGCAGAGGCAGAAGCCCAGGCTTGTTCCACTGAGAAGCACAGGGAAGCCATGCAGAG

Figure S2

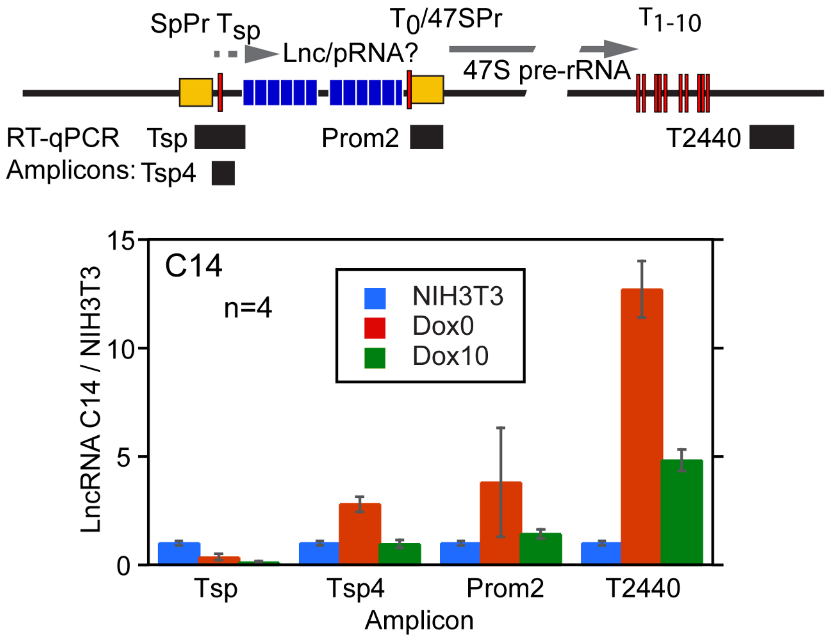
SpPr $T_{sp}$
$T_0$/47SPr
$T_{1-10}$
Lnc/pRNA?
47S pre-rRNA
RT-qPCR
Amplicons:
Tsp
Tsp4
Prom2
T2440
C14
n=4
NIH3T3
Dox0
Dox10
LncRNA C14 / NIH3T3
15
10
5
0
Tsp
Tsp4
Prom2
T2440
Amplicon

Figure S3

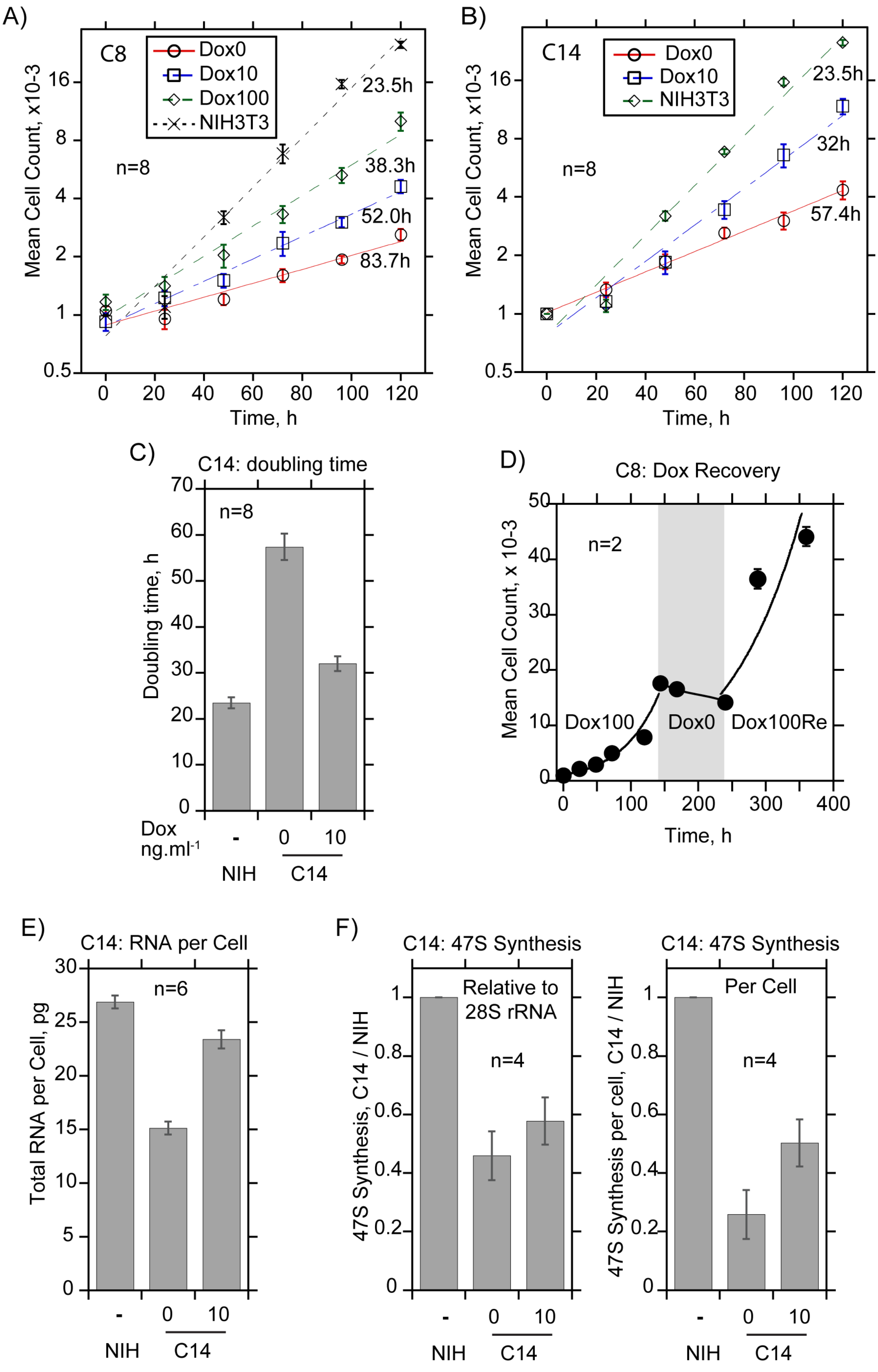
A)
C8
Dox0
Dox10
Dox100
NIH3T3
n=8
23.5h
38.3h
52.0h
83.7h
Mean Cell Count, x10-3
Time, h
B)
C14
Dox0
Dox10
NIH3T3
n=8
23.5h
32h
57.4h
Mean Cell Count, x10-3
Time, h
C)
C14: doubling time
n=8
Doubling time, h
Dox
ng.ml-1
-
0
10
NIH
C14
D)
C8: Dox Recovery
n=2
Dox100
Dox0
Dox100Re
Mean Cell Count, x 10-3
Time, h
E)
C14: RNA per Cell
n=6
Total RNA per Cell, pg
-
0
10
NIH
C14
F)
C14: 47S Synthesis
Relative to
28S rRNA
n=4
47S Synthesis, C14 / NIH
C14: 47S Synthesis
Per Cell
n=4
47S Synthesis per cell, C14 / NIH
NIH
C14

Figure S4

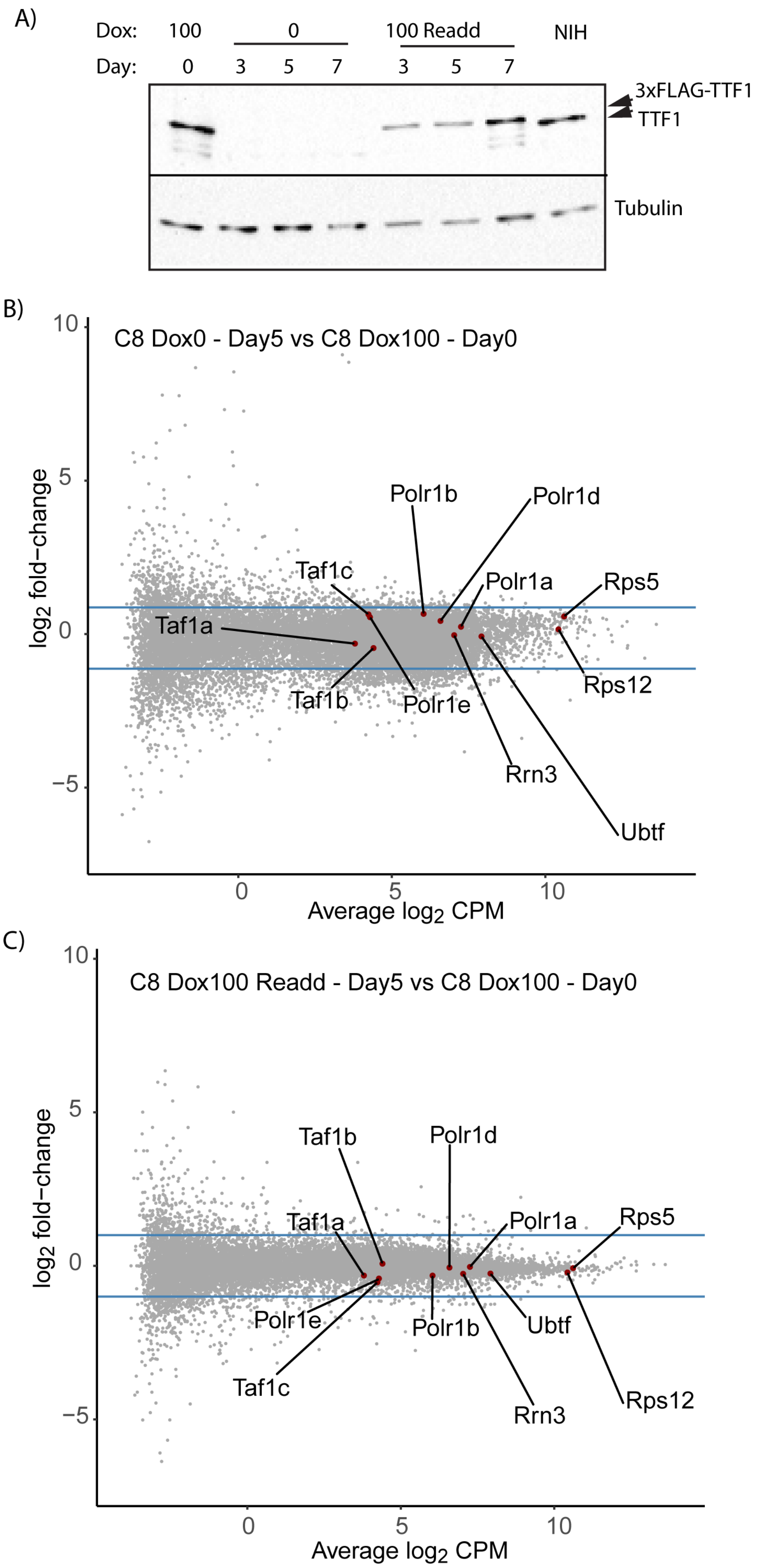
A)
Dox:
100
0
100 Readd
NIH
Day:
0
3
5
7
3
5
7
3xFLAG-TTF1
TTF1
Tubulin
B)
C8 Dox0 - Day5 vs C8 Dox100 - Day0
log2 fold−change
Average log2 CPM
Polr1b
Polr1d
Taf1c
Polr1a
Rps5
Taf1a
Taf1b
Polr1e
Rps12
Rrn3
Ubtf
C)
C8 Dox100 Readd - Day5 vs C8 Dox100 - Day0
log2 fold−change
Average log2 CPM
Taf1b
Polr1d
Taf1a
Polr1a
Rps5
Polr1e
Polr1b
Ubtf
Taf1c
Rrn3
Rps12

Figure S5

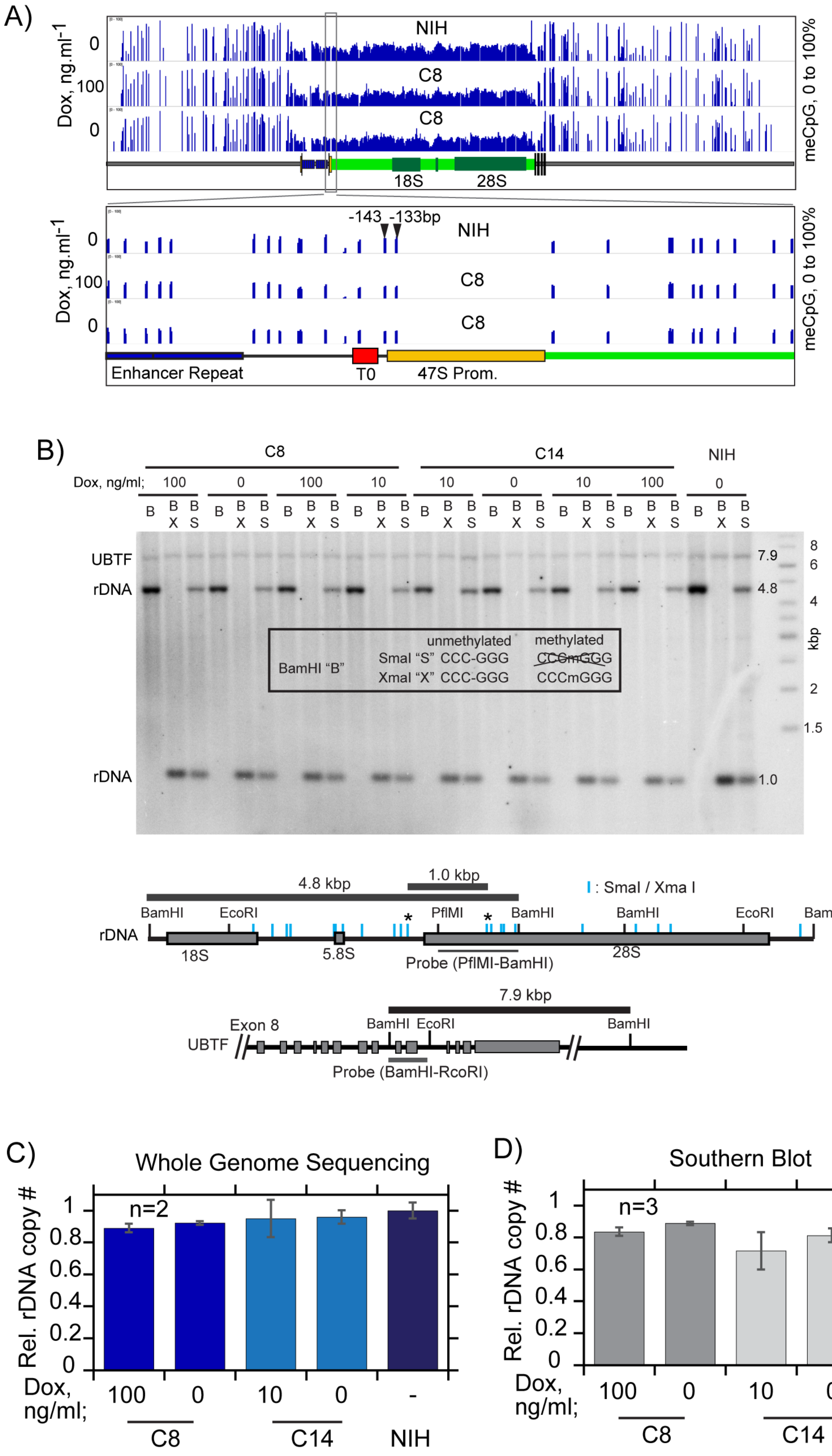
A)
Dox, ng.ml-1
0
100
0
NIH
C8
C8
18S
28S
meCpG, 0 to 100%
-143 -133bp
Enhancer Repeat
T0
47S Prom.
B)
C8
C14
NIH
Dox, ng/ml;
UBTF
rDNA
unmethylated
methylated
BamHI "B"
SmaI "S" CCC-GGG
XmaI "X" CCC-GGG
CCCmGGG
kbp
4.8 kbp
1.0 kbp
SmaI / Xma I
BamHI
EcoRI
PflMI
5.8S
Probe (PflMI-BamHI)
7.9 kbp
Exon 8
Probe (BamHI-RcoRI)
C)
Whole Genome Sequencing
Rel. rDNA copy #
n=2
Dox, ng/ml;
D)
Southern Blot
n=3

Figure S6

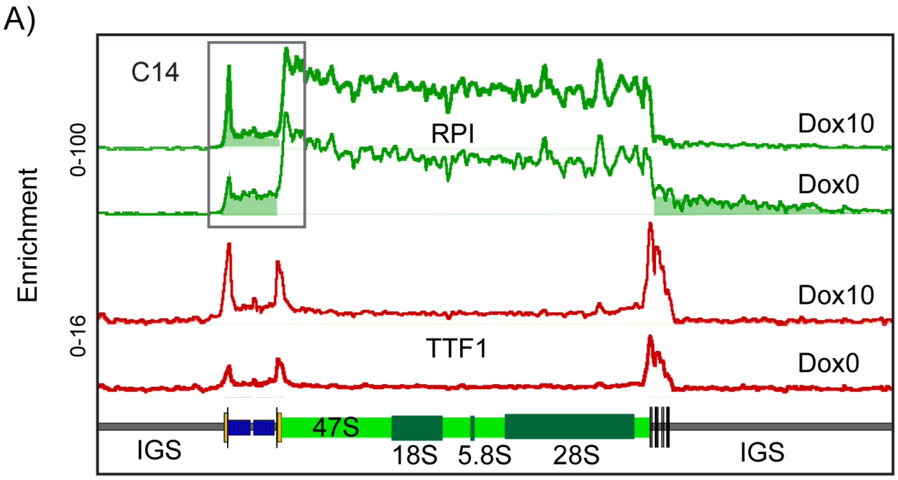
A)
C14
RPI
Dox10
Dox0
0-100
Enrichment
0-16
Dox10
TTF1
Dox0
47S
IGS
18S
5.8S
28S
IGS

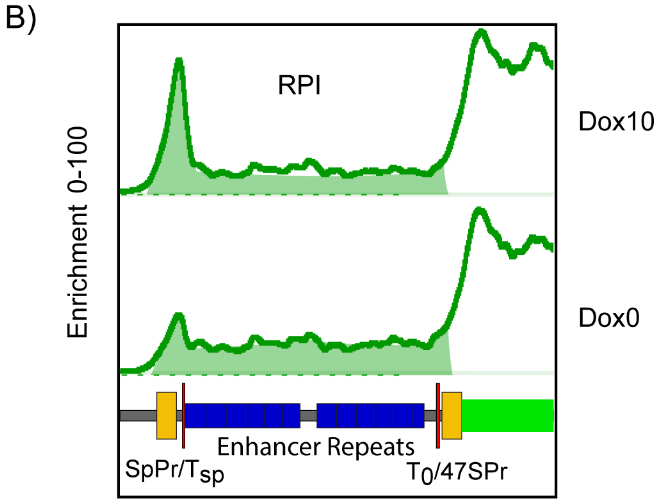
B)
RPI
Dox10
Enrichment 0-100
Dox0
Enhancer Repeats
SpPr/Tsp
T0/47SPr

Figure S7

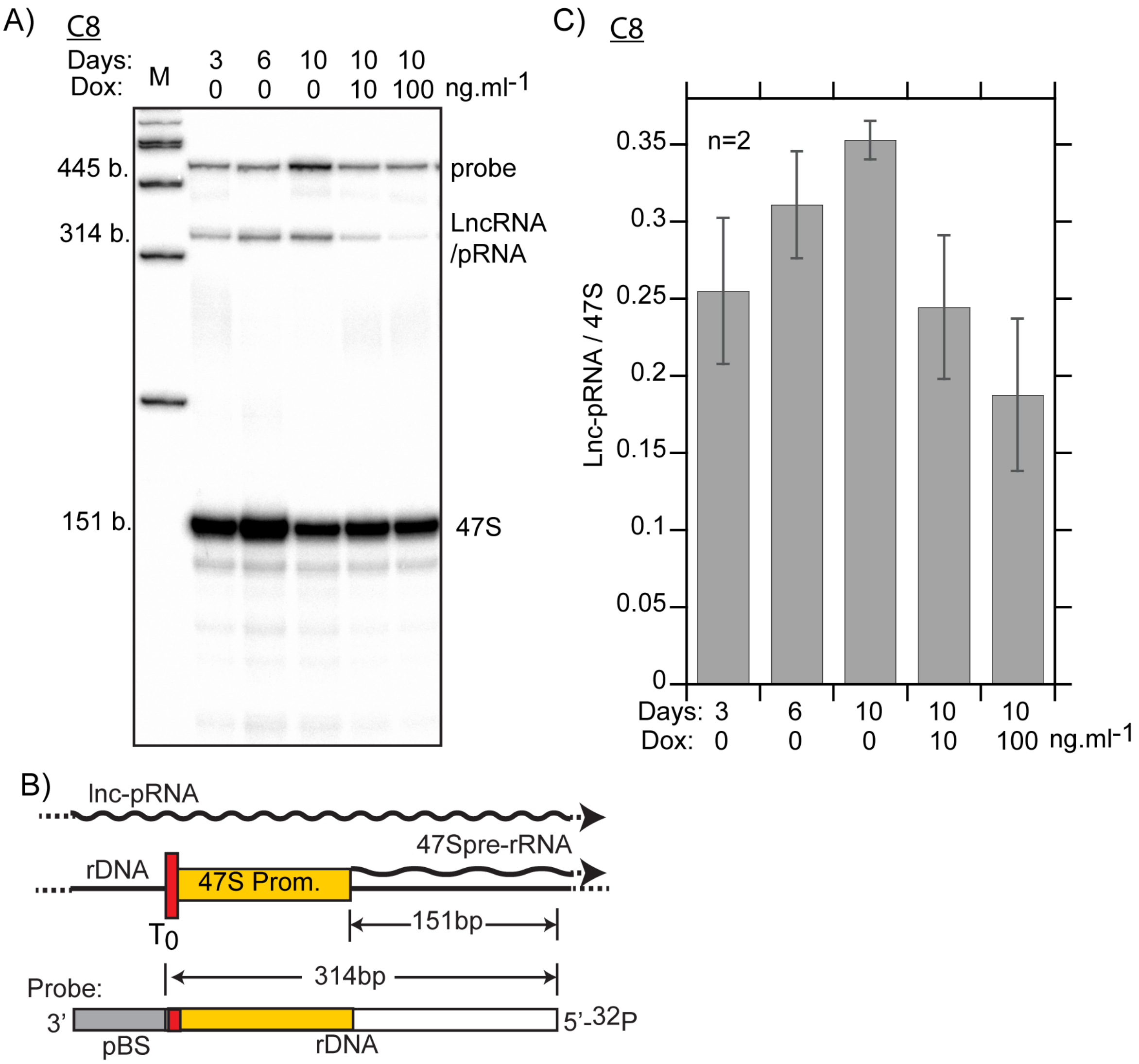
A)
C8
Days: 3 6 10 10 10
Dox: M 0 0 0 10 100 ng.ml-1
445 b.
314 b.
151 b.
probe
LncRNA
/pRNA
47S
B)
lnc-pRNA
47Spre-rRNA
rDNA
47S Prom.
T0
151bp
314bp
Probe:
3'
5'-32P
pBS
rDNA
C)
C8
n=2
Lnc-pRNA / 47S
0.35
0.3
0.25
0.2
0.15
0.1
0.05
0
Days: 3 6 10 10 10
Dox: 0 0 0 10 100 ng.ml-1

Figure S8

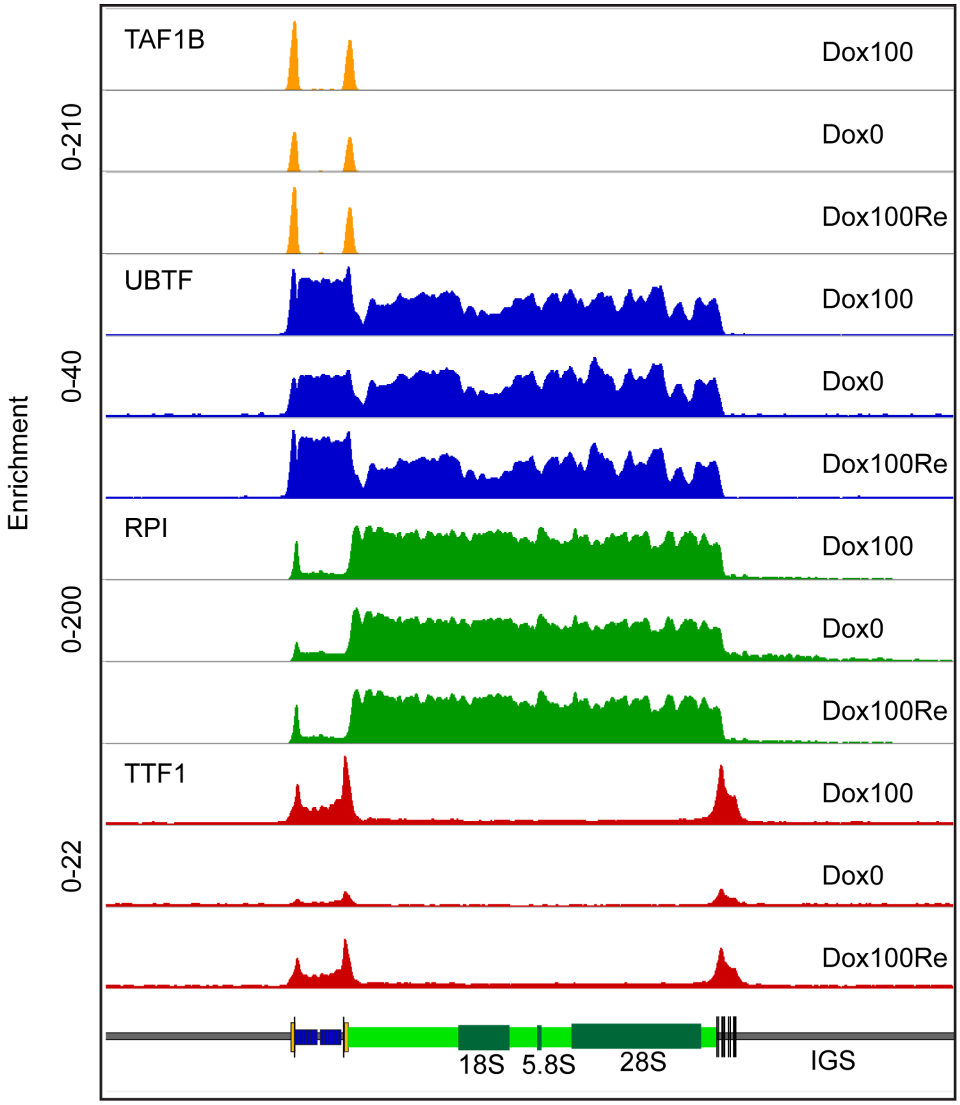
Enrichment
0-210
0-40
0-200
0-22
TAF1B
UBTF
RPI
TTF1
Dox100
Dox0
Dox100Re
Dox100
Dox0
Dox100Re
Dox100
Dox0
Dox100Re
Dox100
Dox0
Dox100Re
18S
5.8S
28S
IGS

Figure S9
